# Odorant Receptor Inhibition is Fundamental to Odor Encoding

**DOI:** 10.1101/760033

**Authors:** Patrick Pfister, Benjamin C. Smith, Barry J. Evans, Jessica H. Brann, Casey Trimmer, Mushhood Sheikh, Randy Arroyave, Gautam Reddy, Hyo-Young Jeong, Daniel A. Raps, Zita Peterlin, Massimo Vergassola, Matthew E. Rogers

## Abstract

Most natural odors are complex mixtures of many volatile components, competing to bind odorant receptors (ORs) expressed in olfactory sensory neurons (OSNs) of the nose. To date surprisingly little is known about how OR antagonism shapes neuronal representations in the periphery of the olfactory system. Here, we investigated its prevalence, the degree to which it disrupts OR ensemble activity, and its conservation across related ORs. Calcium imaging microscopy of dissociated OSNs revealed significant inhibition, often complete attenuation, of responses to indole, a commonly occurring volatile associated with both floral and fecal odors, by a set of 36 tested odorants. To confirm an OR mechanism for the observed inhibition, we performed single-cell transcriptomics on OSNs that exhibited specific response profiles to a diagnostic panel of odorants and identified the receptor Olfr743 which, when tested *in vitro*, recapitulated *ex vivo* responses. We screened ten ORs from the Olfr743 clade with 800 perfumery-related odorants spanning a range of chemical scaffolds and functional groups, over half of which (430) antagonized at least one of the ten ORs. Furthermore, OR activity outcomes were divergent rather than redundant, even for the most closely related paralogs. OR activity fitted a mathematical model of competitive receptor binding and suggests that normalization of OSN ensemble responses to odorant mixtures is the rule rather than the exception. In summary, we observed OR antagonism, inverse agonism and partial agonism occurring frequently and in a combinatorial manner. Thus, extensive receptor-mediated computation of mixture information appears to occur in the olfactory epithelium prior to transmission of odor information to the olfactory bulb.

## INTRODUCTION

An organism’s survival depends, in part, upon its ability to detect and respond to dynamic environments to locate valuable resources, avoid danger, or evaluate conspecific social cues. On land, volatile odorants represent one particularly rich source of information about environmental context, but understanding how this information is detected and processed is challenging. First, estimates of the actual number of odors humans can distinguish are controversial, and have so far resisted reliable empirical determination [1-5]. Compounding this, the chemical diversity of odorant space is extensive and difficult to quantify [6-9], and the complexity of natural odors is also typically high. The same source can exhibit a range of compositions, with component concentrations varying considerably [10, 11]. Additionally, the same component can occur in diverse sources. An example of this is indole, used as a target of this study, which occurs in both fecal [12] and floral sources - it can elicit perceptual qualities reminiscent of both and is widely used in perfumery [13].

In terrestrial mammals (including humans), the task of parsing this stimulus complexity is initiated at the periphery of the olfactory system. Volatile compounds interact with between ∼200-2000 odorant receptors (ORs), depending on the mammal, which are expressed monogenically and monoallelically in olfactory sensory neurons (OSNs) lying in the nasal cavity [4, 14-16]. Demonstration of the combinatorial nature of odor encoding [17-20] appeared to explain how such diverse stimuli could be detected and discriminated by the olfactory system. That is, a single OR can be activated by a range of odorants, and a single odorant can activate multiple ORs [21-28], potentially leading to unique combinations of active ORs for a large number of possible stimuli. However, even with combinatorial encoding and a large family of receptors, recent theoretical work has suggested that OR antagonism may be required to prevent saturation of the peripheral olfactory system and retain discriminatory ability even for modestly complex odors (>20 components) [29].

Since ORs are a large family of class A G protein-coupled receptors (GPCRs), and classic pharmacological experiments on other class A members have revealed disruption of agonist activity via competitive antagonism at the orthosteric site [30-32], it seems likely that ORs would be similarly affected. Indeed, OR antagonism by odorants has been observed experimentally for a relatively small number of mammalian ORs [33-38], and has been projected to play an important role in how odorant mixtures may be encoded [39, 40]. Yet limitations in functional heterologous expression systems and the lack of a protein structure [41] have impeded systematic exploration of the full range of possible receptor-ligand interactions for ORs. As such, it is not well understood to what extent classical GPCR interactions, like antagonism, partial agonism and inverse agonism shape odor representations before they are propagated to the olfactory bulb.

This lack of information on the prevalence of non-agonist interactions, and on correlations between ORs severely limits our ability to understand fundamental aspects of mixture encoding by the olfactory system. For example, infrequent, but strong antagonism would simply prevent the activation of a limited number of ORs in the presence of an antagonist. Alternatively, widespread antagonism could lead to extensive normalization of inputs [29] and decoupling of similarity between mixture compositions and their resulting OSN activity ensembles. With the rise of mathematical models of olfaction demonstrating wide varieties of possible encoding strategies, it has become important to provide data which can help test parameter values and rule out certain classes of model.

In this study, we investigated the prevalence of antagonism, quantified its ability to disrupt OSN activity, and examined its conservation across paralogous ORs using a combination of *ex vivo* calcium imaging microscopy, single-cell transcriptomics and high-throughput *in vitro* OR screening. Here we report evidence that competitive antagonism is a common and fundamental part of odor encoding.

## RESULTS

### Inhibition of olfactory sensory neurons was prevalent in two-component odorant mixtures

The effects of antagonism on the physiological responses of representative populations of OSNs were studied using ratiometric Fura-2 Ca^2+^ imaging of dissociated mouse OSNs. To characterize the reliability of the assay system and precision of odorant administration, dose-dependence and odor response variability to a reference odorant, indole, were established (Figure S1, Methods). OSNs were stimulated with a range of indole concentrations and showed the expected dose-dependent responses and a distribution of sensitivities (Figure S1). The mean EC_50_ across 441 indole-activated OSNs (of 14,989 OSNs, N = 3 experiments) was 54 µM (R^2^ = 0.75, 95% CI [48, 64]). The relative variation in peak height across OSNs repeatedly stimulated with 25 µM indole (corresponding to approximately the mean EC_35_) was 0.79% ± 4.9% (mean ± SD; Figure S1E-F; 86 indole^+^ of 17,412 OSNs, N = 3 experiments).

OSN response modulation was investigated using an experimental design in which the reference odorant was repeatedly delivered (3-4) times and interspersed with delivery of two-component mixtures of the reference odorant and test odorants. This is illustrated in Figure S2 in which the reference odorant was indole at 25 µM and the test odorant was Lilyflore® at 125 µM, a compound which showed a moderate overall level of inhibition compared to the other odorants tested. The peak Ca^2+^ signal evoked by the mixture was compared to the expected peak Ca^2+^ signal of the reference odorant (estimated with a line of best fit across the reference-only stimulus events) allowing for the measurement of relative response changes. Due to the high reproducibility of responses to the reference odorant, any reduction in peak Ca^2+^ response larger than 10% with a test odorant was very unlikely to be due to noise (i.e. >99% of OSNs fluctuated by less than this in the control experiments) (Figure S1D).

Thirty-six odorants were tested for their modulatory effects on indole-responsive OSNs (Figure 1). Previous work has suggested that antagonists for an OR can be structurally similar to its agonists [33, 36-38]; informatively, this appeared to be true in some cases, but by no means all. All tested odorants (Figure 1) revealed reproducible inhibition of greater than 10% in at least 1% of the neuronal populations tested. Furthermore, 17 odorants displayed a median inhibition value greater than 10%. That is, for almost half of the odorants tested, 50% of indole-responsive OSNs showed levels of inhibition greater than 3 SDs above the level attributable to noise. Single-neuron inhibition values up to 100% (complete attenuation) were observed in many OSNs.

**Figure 1.**
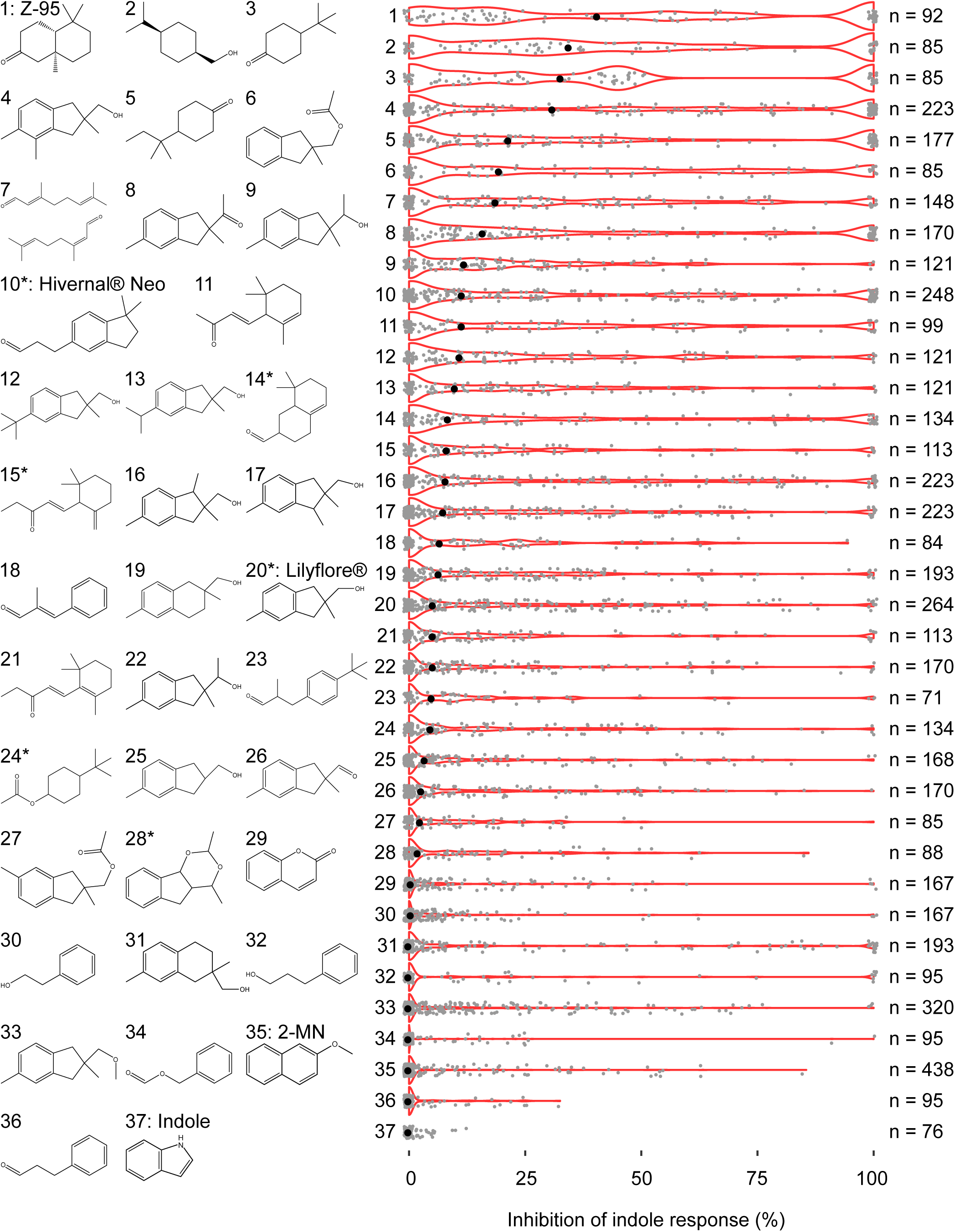
Indole-activated OSNs were inhibited by a range of odorants. Dissociated OSNs were presented with 25 µM of indole alone, or in combination with, 125 µM of compounds 1-36. Violin plots show the level of indole response inhibition where gray dots represent modulation of a single OSN for that odorant, and the black dot denotes the median level of inhibition for the population sampled. The outer red line of the violin repres_3_e_0_nts the probability density at a given inhibition value. 2-MN, 2-methoxynaphthalene. Asterisks denote compounds were tested as a mixture of isomers

### OSN response inhibition was dose-dependent and combinatorial

Previous work has suggested the existence of non-specific CNG-channel inhibition by some odorants [42-45] and other possible non-OR mechanisms could be postulated to explain OSN response inhibition; here we show that this is unlikely for the inhibition that we observed. One characteristic pharmacological property of GPCR antagonism is a sigmoidal dose-dependence spanning several orders of magnitude in concentration. To test for this we chose three inhibitors of indole-responsive OSNs and tested concentrations between 2.5 and 500 µM in mixture with 25 µM indole. The three odorants were chosen to span a range of population effect sizes based on the screening results at 125 µM; Lilyflore® (compound #20) showed the smallest effect of the three with a median inhibition of ∼5%, Hivernal® Neo (compound #10) had a median inhibition of ∼12% and Z-95 (compound #1) showed the greatest effect with a median inhibition level of ∼45%. In all cases dose-dependent inhibition was observed at the single-cell level (Figure 2). Neurons displayed a wide range of sensitivities to the antagonist, such that increasing doses also inhibited the responses of successively larger numbers of OSNs with increasing strength.

**Figure 2.**
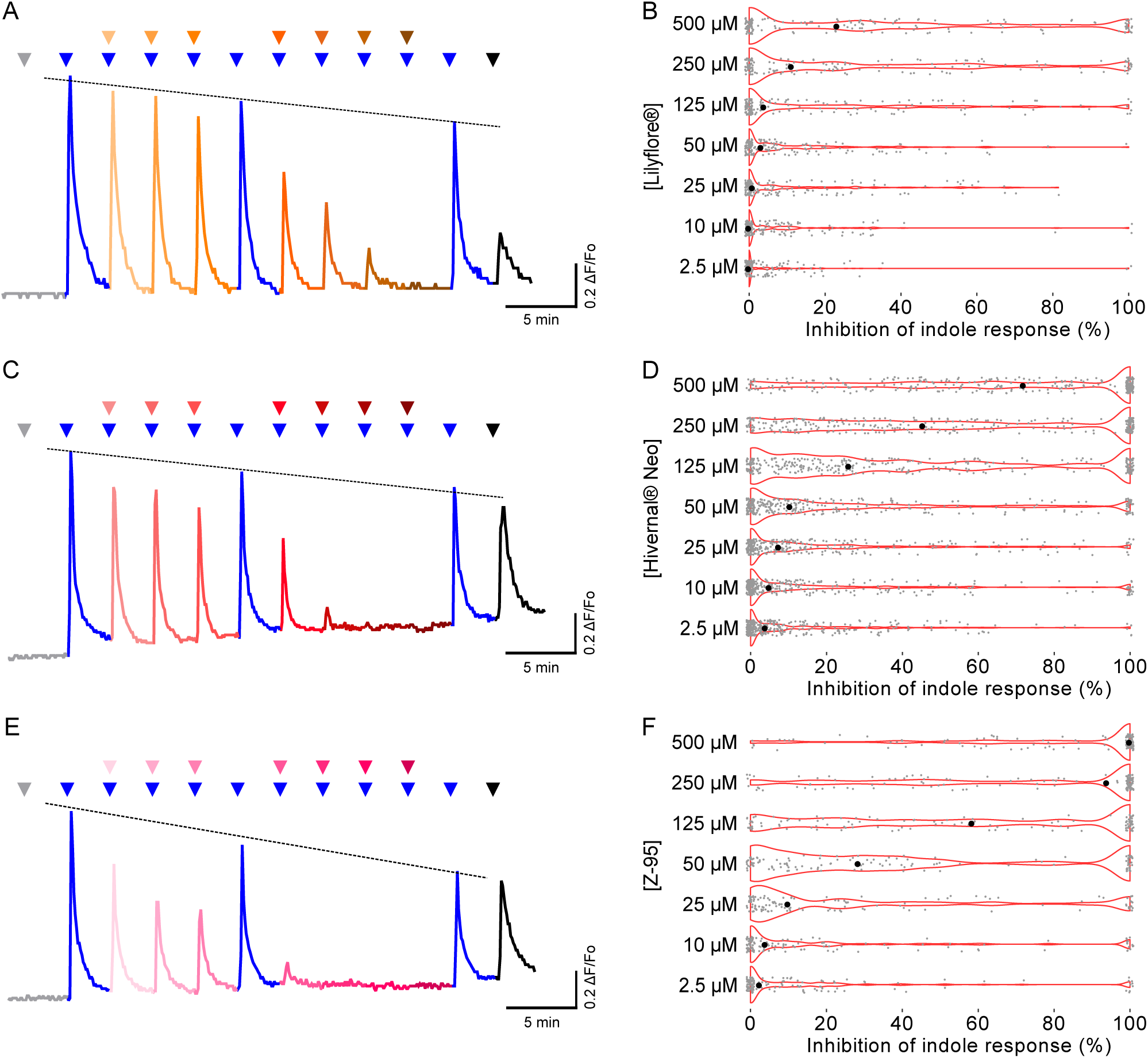
Lilyflore^®^, Hivernal^®^ Neo, and Z-95 inhibited indole-activated OSNs in a dose-dependent manner. (A, C, E) Examples of the dose-dependent inhibition of the indole response by Lilyflore^®^ (A), Hivernal^®^ Neo (C), and Z-95 (E) in separate dissociated OSNs when 25 µM indole alone (in triplicate) or in the presence of increasing concentrations of each inhibitor (in order as shown: 2.5, 10, 25, 50, 125, 250, 500 µM) were administered. Triangles denote the time of administration (gray, 0.3% DMSO, negative control; blue, indole; orange, Lilyflore^®^; maroon, Hivernal^®^ Neo; pink, Z-95; black, 40 µM Forskolin, positive control). (B, D, F) Violin plots showing the inhibition distribution of indole-activated OSNs (25 µM) by increasing concentrations of Lilyflore^®^ (B), Hivernal^®^ Neo (D), and Z-95 (F). Each gray dot represents the modulation observed in a single indole-responsive OSN and the black circle denotes the median value, while the outer red line of the violin represents the probability density at a given inhibition value (for Lilyflore^®^, 115 OSNs, N = 2), Hivernal^®^ Neo (311 OSNs, N = 9), and Z-95 (122 OSNs, N = 3).

While necessary for implicating an OR mechanism, the observed sigmoidal dose-dependence was not sufficient alone to eliminate the possibility of non-specificity. To address this further, indole-responsive OSNs were segregated into subpopulations based on apparent response profiles across a small panel of related odorants. In total, 17 subpopulations of indole-responsive ORs were identified that had differing responses to the panel (Figure 3 and S3), and thus likely represented largely distinct subsets of ORs [33]. Specificity of antagonism was determined by measuring inhibition of the three inhibitors (Z-95, Hivernal® Neo and Lilyflore®) on the same segregated indole-responsive OSNs. When tested, each putative OR antagonist displayed odorant-dependent differences in strength and breadth of inhibition in each sub-population (Figure 3 and S3).

**Figure 3.**
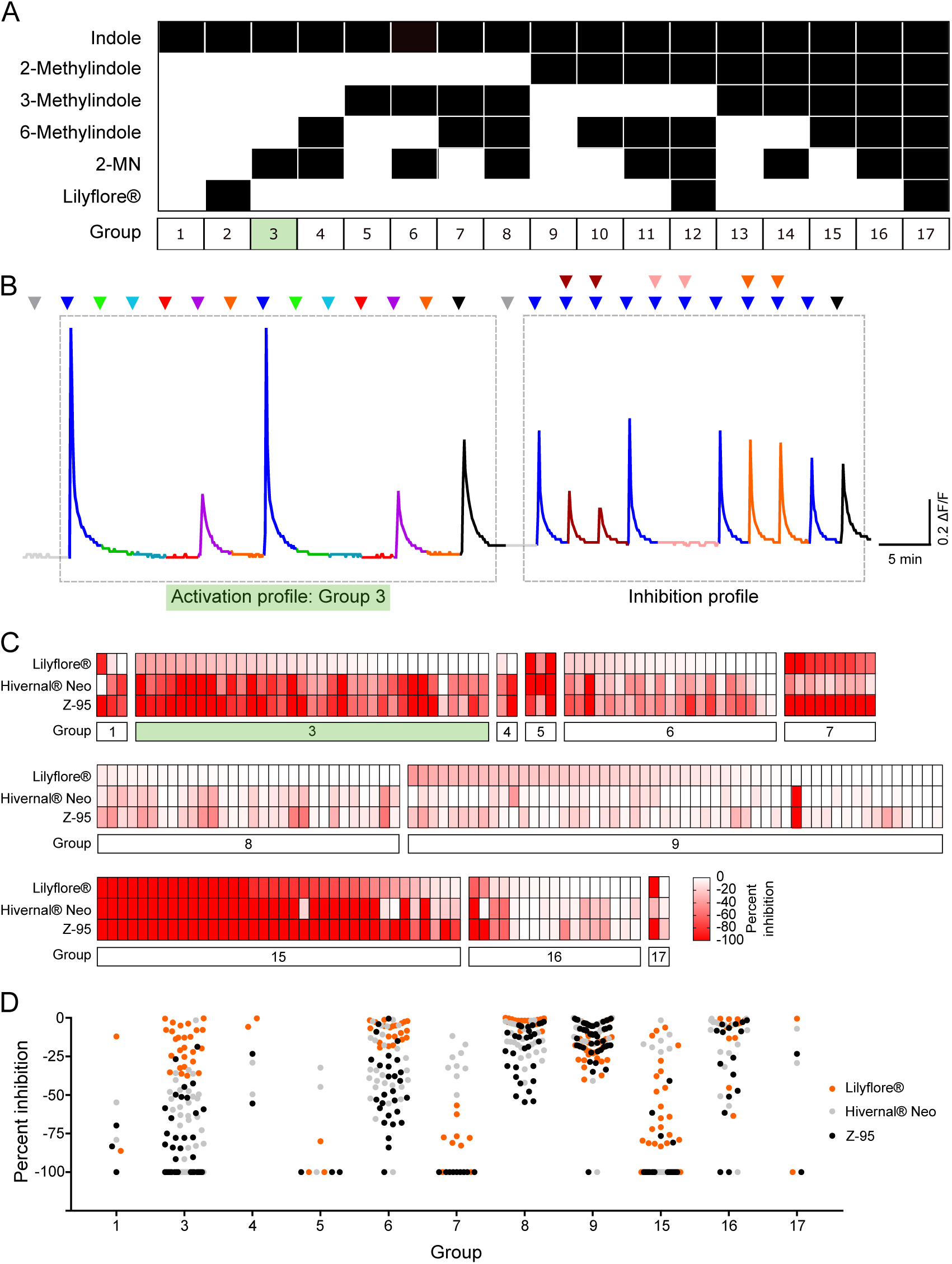
Functionally defined groups of Indole-activated OSNs were differentially inhibited. (A) Binary heatmap of the 17 indole activation profiles that define each group. Group assignment is shown below. (B) Example of an OSN response to a series of singly-administered odorants presented in duplicate (50 µM), including indole (blue), 2-methylindole (green), 3-methylindole (light blue), 6-methylindole (red), 2-methoxynaphthalene (2-MN; purple), and Lilyflore^®^ (orange). Following this series to establish activation profile and corresponding group assignment, indole-activated OSNs were challenged with indole alone (25 µM) or in combination with one of three different inhibitors (125 µM). Triangles denote the time of administration (gray, 0.3% DMSO, negative control; blue, indole; black, 40 µM Forskolin, positive control). The example shown is a group 3 indole-activated OSN by virtue of its co-activation by 2-MN, and is inhibited moderately by Hivernal^®^ Neo (maroon) and strongly by Z-95 (pink), but is not inhibited by Lilyflore^®^ (orange). (C) Population heatmap of the degree of modulation observed by each inhibitor (Lilyflore^®^, Hivernal^®^ Neo, and Z-95) per indole-responsive cell, displayed according to OSN group. (D) Inhibition of indole-activated OSNs by 125 µM of Lilyflore^®^ (orange), Hivernal^®^ Neo (gray), or Z-95 (black) per indole-responsive group described in A.

### Identification of Olfr743, an indole-sensitive odorant receptor

To directly link OSN inhibition to antagonism of specific olfactory receptors, we sought to identify and characterize mouse ORs conferring the observed OSN indole responses. We recorded individual OSNs responsive to sequential stimulation with 50 µM indole and several indole-related compounds (3-methylindole, 2-methoxynaphthalene, 6-methylindole, *para*-cresol; Figure 4A-B). We then isolated cells with high sensitivities to indole for subsequent transcriptome analyses using a single-cell RNA-seq approach. From these analyses several phylogenetically related mouse ORs (Olfr743, Olfr740 and Olfr746) were identified independently. For example, Olfr743 was expressed in two OSNs with similar response profiles (Figure 4 A-B). In all isolated single cells, a single OR transcript was detected at high expression levels, consistent with monogenic OR expression in OSNs [14-16, 18] (Figure 4C-D). The expression of olfactory specific markers, including *OMP, G*_*olf*_, *Adcy3, Cnga2* and *Rtp1*, confirmed each cell was a mature OSN (Figure 4C and 4D insets).

**Figure 4.**
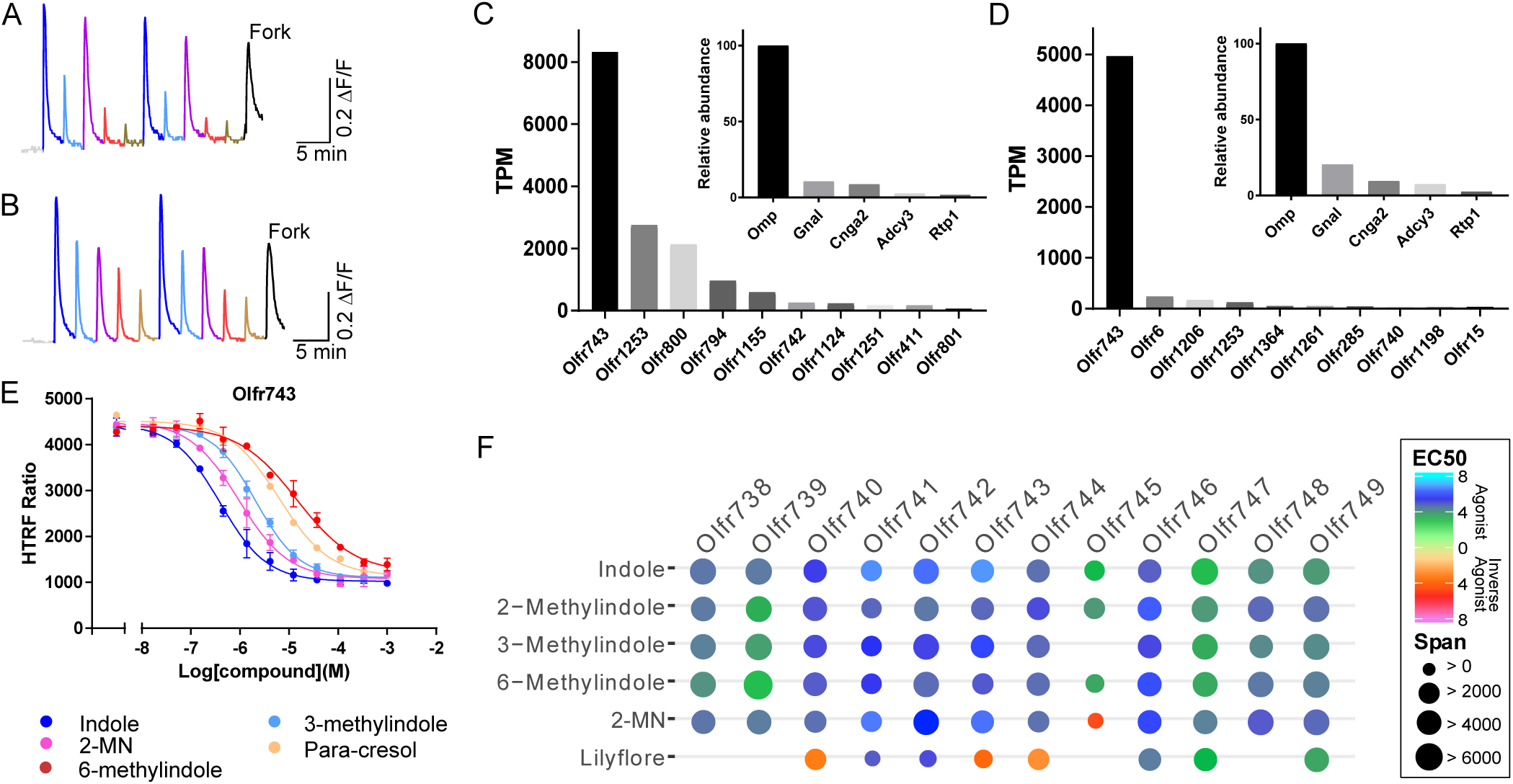
Odorant receptor identification from functionally characterized individual OSNs. (A-B) Ca2+ imaging recordings of two OSNs responding sequentially to 50 µM of indole, 3-methylindole, 2-methoxynaphthalene (2-MN), 6-methylindole and p-cresol. Odorant presentation scheme was repeated twice for consistency and OSN viability assessed by 40 µM Forskolin (Fork). (C-D) Single-cell RNA-seq results corresponding to the cells shown in A and B respectively. OR transcriptome analysis of both isolated OSNs revealed the expression Olfr743 as the most abundant odorant receptor mRNA. Inset shows the expression of Olfactory Marker Protein (Omp), Golf (GnaI), Cyclic Nucleotide-Gated Channel A2 (Cnga2), Adenylyl Cyclase 3 (Adcy3) and Receptor Transport Protein 1 (Rtp1), scaled to OMP, and confirmed the olfactory and neuronal origin of the isolated cells. X axis: RefSeq gene names. Y axis: Transcript Per Million (TPM) reads. (E) Response profile of Olfr743 to increasing doses of indole-related ligands used for the OSN characterization. Due to the competition-based nature of the HTRF assay, a decreasing HTRF ratio indicates cAMP accumulation and a corresponding activity increase. Both the potency and efficacy of Olfr743’s response to these ligands closely matched the OSN activity ranking as evaluated by peak height in A and B. X axis, log concentration in Molarity. Y axis, activity relative to cAMP HTRF ratio. (F) Olfr743-related ORs (see Figure S4) were tested at a range of concentrations with a diagnostic panel of ligands. In the resulting heat map blue and red hue represent the calculated EC50 and IC50 for agonism and inverse agonism, respectively. Activation efficacy is represented by the diameter of the circle. Absence of a circle indicates absence of activation or inverse agonism but does not exclude neutral antagonism.

We then characterized the activation profile of Olfr743 by performing dose-response experiments in a functional heterologous expression system with the same compounds used to characterize the OSNs of origin. We used an immuno-competitive cAMP binding assay and measured the receptor’s specificity (the molecular receptive range within the confines of our library), and odorants’ potency (sensitivity to a particular ligand or EC_50_) and efficacy (the extent of activation between no binding and full saturation of the receptor). With this approach, the recorded signal is inversely proportional to the concentration of cAMP produced upon ligand binding. The response of Olfr743 to these compounds varied in potency and in efficacy (maximum HTRF ratio) (Figure 4E). Importantly, the compound response rank-order was largely consistent with that of peak heights (used as a proxy for OSN sensitivity) in the OSN responses. Overall, these results suggest that Olfr743 is directly responsible for the observed OSN responses, and provide an effective method to deorphan ORs based on complex response profiles [46].

### Characterization of an indole-responsive odorant receptor gene family

Because distinct OSN response profiles indicated that more than one OR could respond to indole (Figures 3 and S3), we extended the search for indole-sensitive ORs to a seven-member paralogous gene family with greater than 78% amino acid identity to Olfr743. In addition to this first group, we also selected an additional set of five receptor genes with greater than 55% identity, defining a less conserved phylogenetic outgroup (Figure S4). We first tested the response of the 12 receptors to increasing doses of indole and structurally-related compounds (Figures 4F and S5). All paralogs except Olfr745 were activated by all of the indole-related odorants except Lilyflore®. Although the receptors showed similar and partially overlapping activation profiles, potency varied widely among receptors for a given ligand, in some cases spanning several orders of magnitude. For example, indole EC_50_s obtained from complete dose-response curves ranged between nanomolar and micromolar concentrations (80 nM for Olfr743 to 81 µM for Olfr748). Even among paralogous ORs sharing almost 80% identity, indole’s EC_50_ still varied by more than two orders of magnitude (from 80 nM for Olfr743 to 32 µM for Olfr739). Response efficacy also varied, encompassing inverse, partial and full agonism. 2-methoxynaphthalene partially activated Olfr738, Olfr739, Olfr740, and Olfr744, (71-75% of the maximum indole response) and fully activated Olfr741, Olfr742, and Olfr743. Similarly, Lilyflore® displayed no, partial and full agonism across the tested ORs, even eliciting clear inverse agonism on Olfr740, Olfr743 and Olfr744, indicating that the latter three receptors were constitutively active in our assay (Figure S5). Thus, even across this small set of ligands, these closely related ORs displayed diverse agonist profiles, and, moreover, displayed receptor binding outcomes not confined to full agonism; namely partial and inverse agonism.

### Functional diversity of the Olfr743 family

We examined the range of functional responses for this family by performing single-concentration agonist and antagonist screens using ∼800 distinct stimuli for the 10 ORs which displayed a full indole-dose response (Figure S5). These compounds consisted of 640 aromatic and 282 aliphatic compounds, covering 195 combinations of various chemical features (Figure S6). Agonism was assessed by stimulating each OR with 300 µM of each compound, while antagonism was assessed by measuring the change in receptor activation elicited by 300 µM of each compound in the presence of the EC_80_ of indole for each OR. We then chose the top 5 antagonists or agonists for each OR and tested these compounds against all 10 ORs in dose-response experiments in the presence (for antagonists) or absence (for agonists) of the EC_80_ of indole. Using these data, we constructed receiver operating characteristic (ROC) curves for each screen. The area under the curve (AUC) for the antagonist screen was 0.78, p < 0.0001; AUC for the agonist screen was 0.90, p < 0.0001; (Figure S7). From this analysis, 75% of true antagonists showed inhibition greater than 24% in single-concentration antagonist screens and 80% of true agonists showed activation greater than 32% in single-concentration agonist screens.

Using these cutoff values, we then examined the number of antagonists and agonists for each OR and the overlap among related family members. We identified a total of 430 antagonists and 444 agonists that elicited a response in at least one OR, i.e. over half the tested library for both screens (Figure 5 and S8). The number of antagonists and agonists varied widely and tended to be inversely correlated. For example, Olfr746 had 30 antagonists and over 400 agonists, while Olfr740 had the greatest number of antagonists (352) and the lowest number of agonists (28) (Figure 5A). We used an Upset plot [47] to examine the number of unique antagonists and agonists for each OR as well as the overlap among ORs (Figures 5B and S8A). Of the 430 antagonists, Olfr740 had both the largest number of total and unique antagonists (149). Olfr746 had both the largest number of total and unique agonists (187).

**Figure 5.**
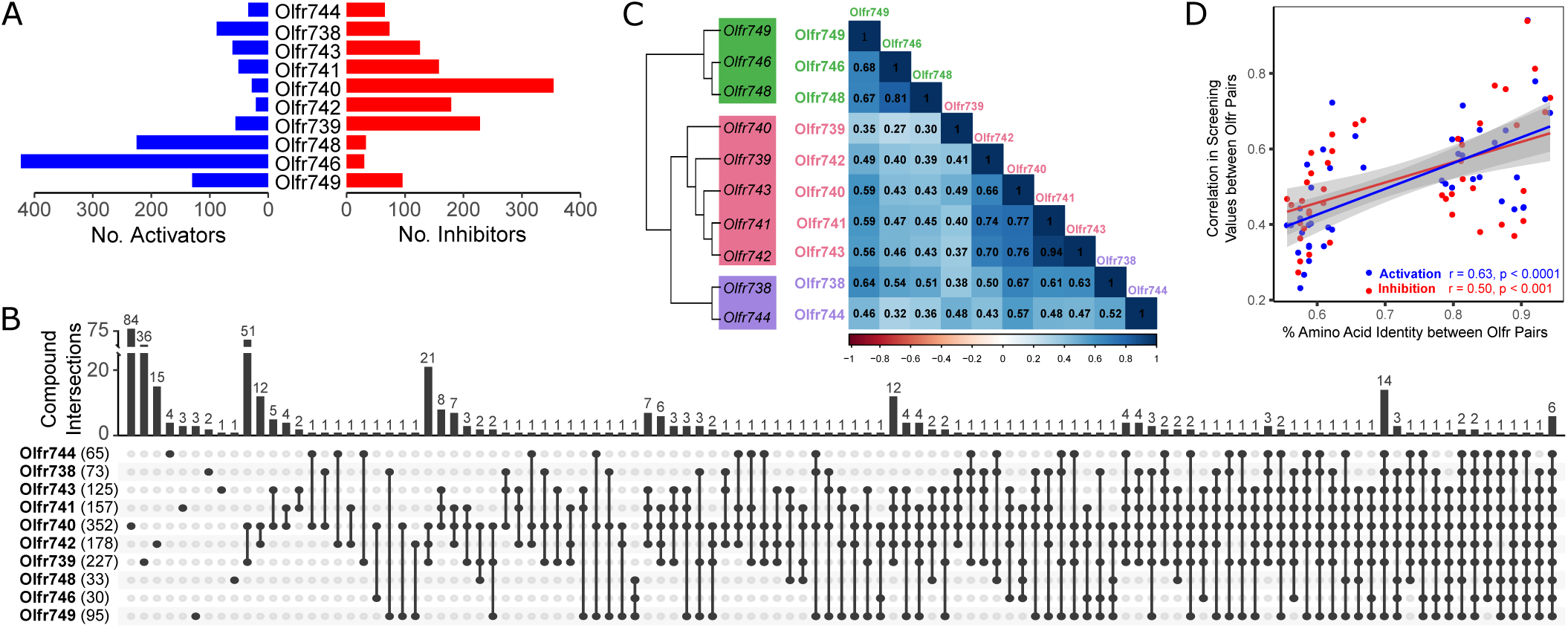
Large inhibition screen reveals antagonism diversity among closely related indole-sensitive ORs. (A) The number of activators (> 32% activation) and inhibitors (> 24% inhibition) identified from single-point concentration screening of 700 (activation screen) or 800 (inhibition screen) compounds against the Olfr743 family. (B) An upset plot showing the number of inhibitors (bar height) which were unique (single dots) or shared (linked dots) among Olfr743 family members. (C) Cladogram representing the phylogenetic relationship between the ten receptors screened (based on amino acid alignment) is shown in parallel to the pairwise Pearson correlation matrix obtained from the full modulation screening results (all points considered). (D) Correlation between the percent amino acid identity and correlation in either inhibition (red points) or activation (blue points) screening values for each OR pair.

Many of the compounds tested were ligands for multiple Olfr743 family members – 281 were antagonists, and 257 agonists for more than one of the tested ORs. For OR pairs with greater than 90% identity, the average frequency of overlap was 0.70 for inhibitors and 0.86 for activators, yet we observed variation in this frequency. For some closely-related pairs of receptors, the number of overlapping compounds reflected their degree of relatedness. For example, Olfr741 and Olfr742, which have 94% amino acid identity, shared 75% of their antagonists and 100% of their agonists (i.e. all Olfr742 agonists also activated Olfr741). For some receptor pairs, however, the number of antagonists or agonists shared between two ORs was less reflective of their similarity. Olfr739 is 90% identical to Olfr742 and shares only 60% of its antagonists and 67% of its agonists.

To further compare the functional responses of this OR family, we constructed Pearson correlation matrices for the antagonist (Figure 5C) and agonist (Figure S8B) screening values for each pair of ORs. Hierarchical clustering of the correlation matrices recapitulated phylogenetic proximity for both screens; however, the median correlation values were moderate (r_median_ is ∼0.5 for both assays) and varied extensively, indicating high, but widely variable functional diversification (Figure S8C). As shown in Figure 5C, Olfr741 and Olfr742 were highly correlated in both their inhibition and their activation profiles (r_median_ ∼0.70 for both assays), whereas, Olfr739 and Olfr742 were much less well-correlated (r_median_ ∼0.4 for both inhibition and activation) despite sharing 94% and 90% amino acid identities, respectively. In fact, an OR pair with relatively low identity, Olfr738 and Olfr749 (∼60%), was better correlated for both activation (r = 0.72) and inhibition (r = 0.63) than Olfr739 and Olfr742. When compared directly (Figure 5D), the correlation between percent amino acid identity and functional similarity (as measured by the pairwise OR Pearson correlation values) reflected a high degree of functional divergence with statistically significant (p < 0.001 for inhibition and p < 0.0001 for activation), but low positive correlations (r = 0.50 for inhibition, r = 0.63 for activation). Furthermore, when analyzed by calculating Jaccard indices over the sets of ligands (Figure S9, Methods), the agonists and antagonists again showed very similar distributions of values, but were less conserved, on average, than binding was. These results indicate that phylogenetic similarity is not always indicative of functional similarity, and vice versa. Also implied, is that as ORs diversify over evolutionary time, binding outcomes (i.e. agonism and antagonism) over a set of odorants diversify more rapidly than the ability to bind them.

### Widespread and diversifying antagonism of related ORs

To further explore the diversity of functional outcomes among these paralogous ORs, we measured their activation (EC_50_), and inhibition dose-response profiles (relative IC_50_ in the presence of a potent concentration of indole, EC_80_), for the top five antagonists for each OR as well as indole and Lilyflore® (Figure 6A). The 37 compounds tested in this way can be seen to span a range of chemical scaffolds, including 5,6- and 6,6-bicyclic aromatics, macrocycles and polycycles, aliphatic chains, and multiple functional group including alcohols, ketones, acids, lactones and esters. A composite heat map including both measurements summarizes the diversity of outcomes we observed (Figure 6B). The IC_50_ rank-order of the top five antagonists selected for each receptor was consistent when compared to the single-concentration inhibition modulation values. All 37 compounds inhibited more than one receptor (c.f. Figure 5), consistent with the high frequency of antagonism previously observed. Compounds #1, #10, #11 and #42-45 inhibited all receptors despite sequence identities as low as 56% between Olfr746 and Olfr741 (Figure 6B and S4). As in Figure 4, indole again activated all of these ORs. Almost half of the compounds (16, 44%) showed divergent outcomes at the concentrations tested, ranging from strong inhibition to potent activation. This was also observed for the closest paralogs. Olfr741 and Olfr742, with 94% sequence identity and greater than 0.70 correlation in antagonist and agonist responses, showed qualitatively different outcomes for six compounds (#47 and #59 inhibiting only Olfr742; #52, #56, #63 displaying opposite outcomes). Similarly, Olfr741 and Olfr743, which displayed the highest correlation in antagonist and agonist response (r = 0.94 for both assays), showed a divergent response to six compounds (#20, #54 and #64 inhibiting only Olfr742; #53 activating only Olfr741; #52 displaying opposite outcome). Finally, as an example of a receptor pair that is highly genetically similar but not functionally so, Olfr742 and Olfr739 differed in their response to nine of the tested compounds. As with single-concentration screening values, hierarchical clustering of similarity in OR response profile across the 37 compounds again largely reflected phylogenetic clustering (Figure 6B, top). However, full dose-response experiments revealed extensive functional divergence even in cases where correlations between single-concentration screening responses indicated a high level of functional similarity. Taken together, these data support the results obtained from the single-concentration screening and their associated implications of widespread, dose-dependent antagonism and its role in functional diversification even among paralogous receptors.

**Figure 6.**
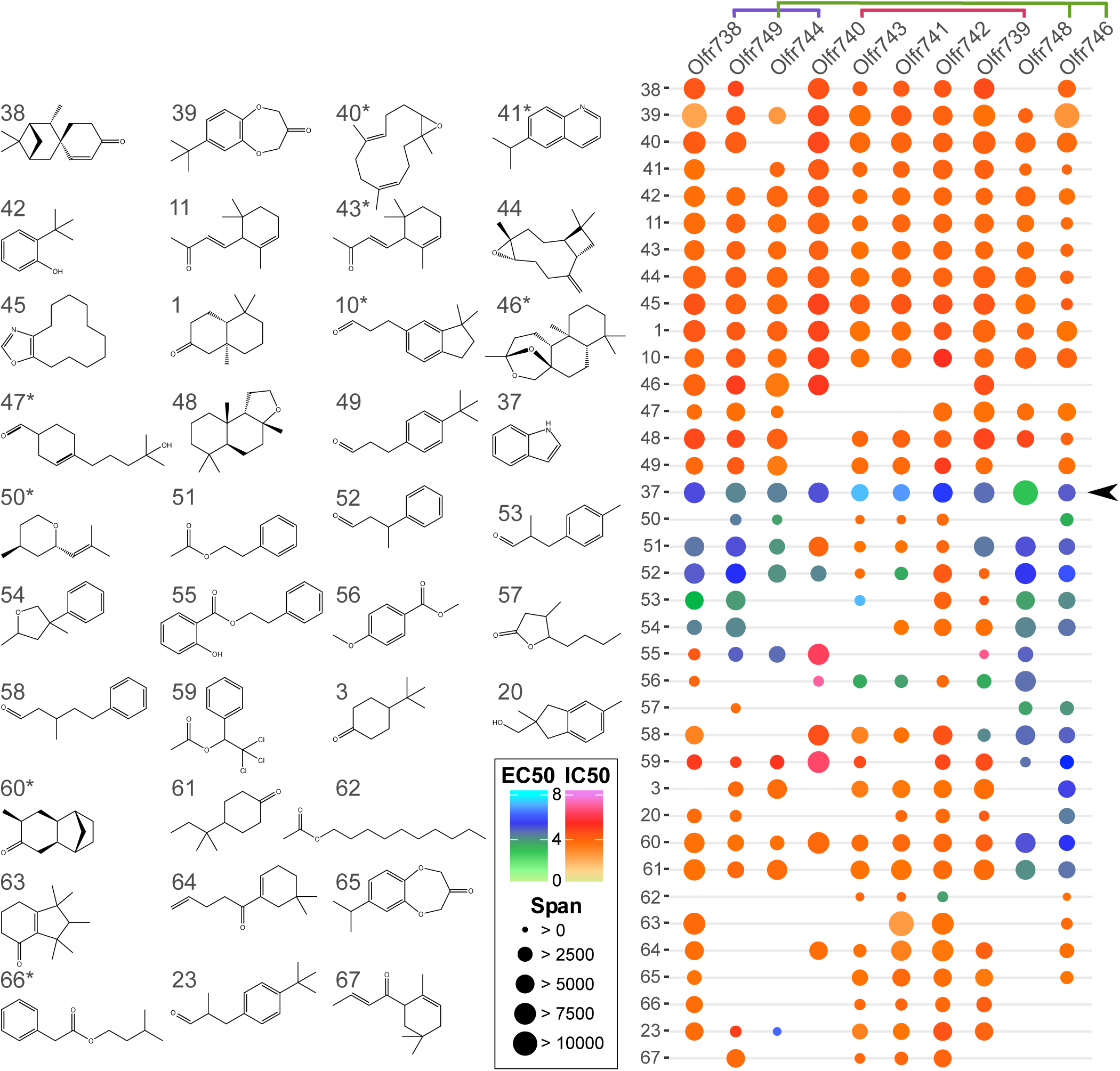
Combinatorial inhibition and activation of Olfr743 family. (A) The 35 strongest antagonists (top five non-redundant hits per OR) identified in the single-concentration screen, as well as indole and Lilyflore^®^ were tested for agonist and antagonist dose-responses. Compounds with an asterisk (^⋆^) are mixtures of isomers (with only one shown, for simplification). (B) Composite heat map summarizing full dose-response curves obtained for both the inhibition (red hue IC50) and activation (blue hue EC50) of the Olfr743 receptor family; no circle indicates the absence of significant ligand binding for a particular OR. When a dose-response curve could be fitted in both inhibition and activation mode (i.e. partial agonists), the IC50 is shown. The functional diversification elicited by exposing phylogenetically-related ORs to binary mixtures is readily visible and includes inhibition, activation and no measurable effect. Potency scale in - log molar concentration.

### Binding affinity is weakly correlated with functional outcome

Mathematical models suggest that the correlation between binding affinity and activation efficacy plays an important role in odor identification and discrimination, assuming mixture interactions are primarily due to competitive binding at the receptor level [29, 48-50]. In short, if the binding affinity and activation efficacy (i.e. ability to induce OR downstream signaling once bound) are largely uncorrelated, agonism and antagonism across the receptor ensemble are balanced. This reduces the chances of saturating the receptor repertoire in the presence of complex odor mixtures and may allow for greatly improved odor segmentation and discrimination.

In order to quantitatively test whether antagonism is indeed mediated through competition for receptor binding between the activator and inhibitor, we performed a competitive binding assay for Olfr740 with indole as the activator and compounds #10 and #1 as the inhibitors, with both agonist and antagonists varied over a range of concentrations. The data was fit to a two-step competitive binding (CB) model (Figure 7A, Methods), where the activator and inhibitor compete to bind to the receptor with binding affinities κ^-1^_act_ and κ^-1^_inh_ respectively. The bound compound then activates the receptor with an activation efficacy η_act_ or η_inh_, depending on whether the activator or inhibitor is bound. The model constrains the activation efficacy to lie between 0 and 1, where 1 represents a perfect agonist i.e., the compound fully activates the receptor whereas 0 represents no activation. To explain the cases of inverse agonism observed in the dataset, we added the possibility for spontaneous activation of the receptor which introduces constitutive activity η_s_ (between 0 and 1) for the receptor when no compound is bound. The key parameter that determines a compound’s functional outcome is its activation efficacy. A compound is a partial agonist if its activation efficacy η is less than 1 but greater than η_s_, and is an inverse agonist if η is less than η_s_. If η_act_ > η_inh_ and c_act_/κ_act_ < c_inh_/κ_inh_, then the inhibitor ‘antagonizes’ the activator. The model yields an excellent fit to the data for the two inhibitors tested (Figure 7B), using three parameters: the binding affinities of the activator and inhibitor, κ^-1^_act_ and κ^-1^_inh_ and the ratio of the subtracted activation efficacies (η_inh -_ η_s_)/(η_act -_ η_s_) (Methods).

**Figure 7.**
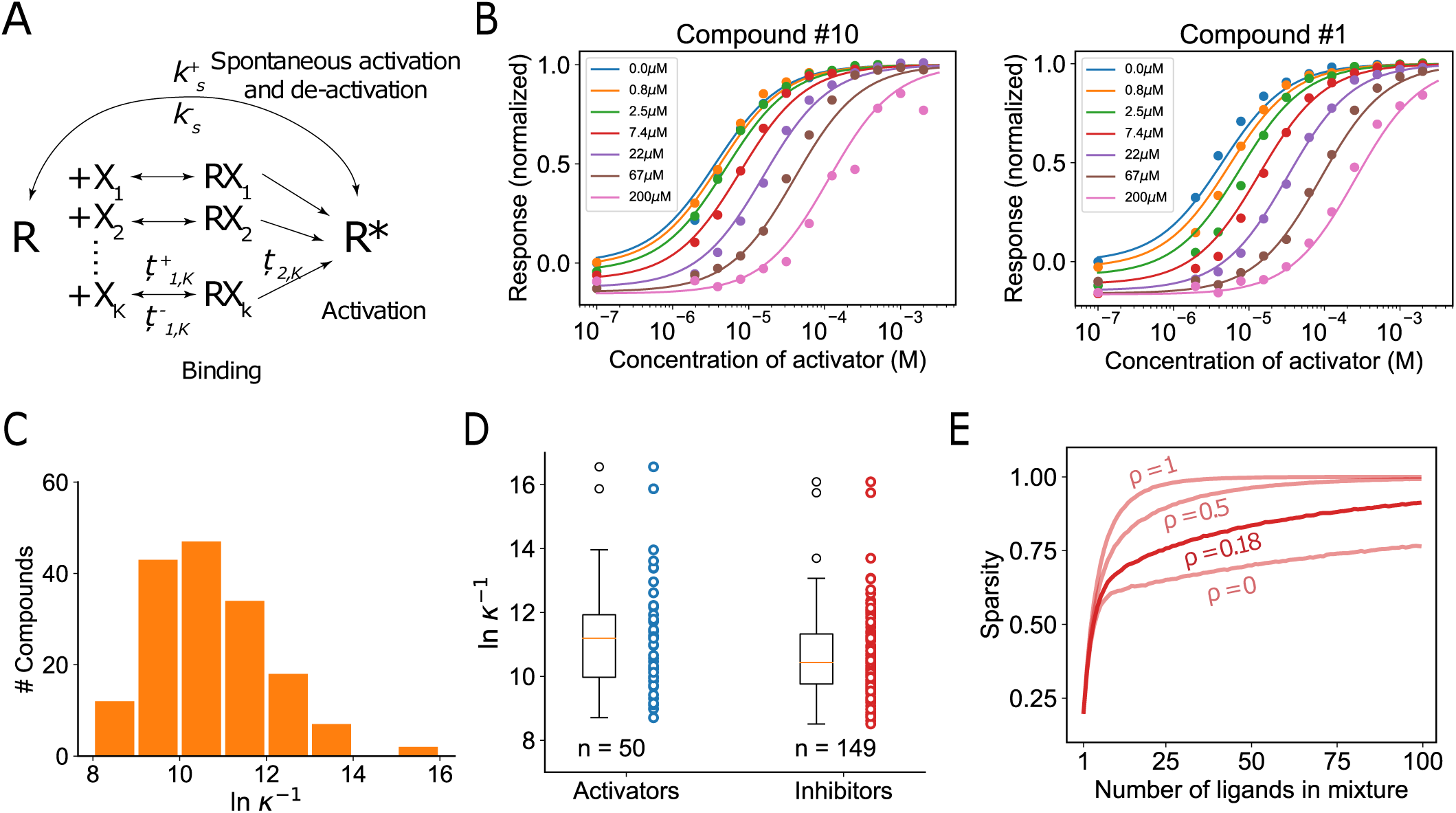
The competitive binding model of mixture interactions and the relationship between binding affinity and functional outcome. (A) A schematic of the competitive binding (CB) model with spontaneous activity. (B) Data from a CB assay using Olfr740 with indole as the activator and compounds #10 and #1 as the inhibitors. Data (solid circles) show the activation profiles for different concentrations of the inhibitors (legend). The solid lines show the best fit curves using the CB model from A. The data is normalized between 0 and 1. The values below 0 indicate suppression of the constitutive activity by the inhibitor. (C) Histogram of binding affinities obtained by fitting the CB model to activation and inhibition profiles for 39 compounds across the Olfr743 receptor family. (D) Box and scatter plots showing the binding affinities for activators and inhibitors. While the distributions are largely overlapping, activators have a slightly higher binding affinity (median = 11.0) compared to inhibitors (median = 10.46) leading to a weak, but non-zero correlation between binding affinity and functional outcome. (E) The probability that a receptor is activated (sparsity) increases slower with mixture complexity when the correlation between binding affinity and functional outcome is weak compared to any additive model of mixture interactions (= 1), and thus prevents saturation of the receptor ensemble.

To measure the correlation between binding affinity and activation efficacy, we fit the competitive binding model to the activation and inhibition profiles of 39 unique compounds for each of the ten ORs in the Olfr743 family, where the activator indole was delivered at EC_80,_ as before. The distribution of ln κ^-1^ (*n* = 199) was found to be close to a normal distribution (mean = 10.76 ∼ 20 μM in concentration, SD = 1.37) with a few sensitive outliers (Figure 7C). We found a weak correlation between the binding affinity of activators (η ≈ 1, *n*_*act*_ = 50) and inhibitors (η ≈ 0, *n*_*inh*_ = 149) of ρ = 0.18 ± 0.03 (mean ± SD), where ρ is the Pearson correlation coefficient between ln κ^1^ and η (Figure 7D). To test whether the measured statistics lead to the receptor activity normalization predicted in theory [29], we numerically computed the extent of saturation of a receptor ensemble with increasing mixture complexity, where mixture interactions were modeled using the competitive binding model and the receptor-ligand binding and activation parameters were drawn from the experimentally measured distributions (Figure 7E). The fraction of activated receptors (sparsity) indeed increased at a slower rate compared to any additive model of mixture interactions (ρ = 1), reaching 90% saturation for > 100 ligands compared to ≈25 for an additive model. Thus, competitive binding appeared to account for the widespread antagonism observed. Binding affinity and activation efficacy appeared largely decoupled (ρ∼0.18) and these statistics would result in a meaningful expansion of the encoding capacity of the system compared to one in which antagonism was rare (ρ→1) (Figure 7E).

## DISCUSSION

Natural odors in the environment are typically mixtures of many different chemicals, yet the precise mechanisms by which complex mixtures are encoded by the odorant receptor repertoire are yet to be elucidated. In particular, the role of OR antagonism in this regard has lacked extensive empirical characterization, which formed the primary focus of the current study. We found evidence for the involvement of the full spectrum of competitive, non-cooperative, GPCR interactions in a way that was not especially biased towards agonism.

Indole was chosen as a target for antagonism throughout the study as an example of an interesting, but presumably not unique, molecule. It occurs in nature, is present at substantial levels in some flowers, and is used regularly in perfumes [13]. It is also present in latrine headspaces [12] and is reminiscent of feces if delivered at high concentrations. We found that indole and other structurally related odorants robustly and dose-dependently activated different subsets of *ex vivo* mouse OSNs. However, one of the ingredients tested (Lilyflore® - Figure S2) showed minimal activation overlap, despite sharing a similar 6,5-bicyclic ring structure. We thus hypothesized that Lilyflore® may still bind indole OSNs, but instead inhibit their activity. Indeed, this was the case not just for Lilyflore®, and additional structurally related compounds, but other common perfumery ingredients with varied chemical structures as well. For many odorants, complete OSN inhibition affected a large subset of the OSN population. Inhibition was much more common than the lack thereof, even considering the relatively small selection of compounds tested.

Given this finding, it was necessary to show that the OSN inhibition was indeed mediated by antagonism of indole-responsive ORs and to systematically explore its prevalence. Sampling >10,000 murine OSNs with up to 30 odorant presentations, we characterized the heterogeneous indole activation code, finding it could be partially deconstructed by examining the activation overlap between indole and five structurally related analogues. Using this panel, we identified 17 specific activation profiles within the indole-responsive OSN population, allowing us to subsequently probe the specificity of OSN response inhibition for three of the inhibitors. While these 17 groups are expected to largely segregate different OSN types (i.e. OSNs each expressing different ORs), it was, however, not possible to use this technique to definitively relate inhibitor specificity to specific OSN types present in the indole-responsive population. This is because group mis-assignment of OSN types can occur due to the concurrent presence of several factors, including variation in sensitivity of OSNs of the same type and the relatively small number of diagnostic ligands used. For example, in comparing the *in vitro* responses of the identified OR family to the *ex vivo* responses, most of the ORs were consistent with only a small subset of the OSN groups (i.e. the majority were consistent with groups 16 and 17, while Olfr745 was most similar to group 10, in Figure 4B). Nonetheless, this approach provided a powerful way to observe the specificity of inhibition by different ligands, which when coupled with their sigmoidal dose-dependence, provided strong evidence for widespread OR antagonism.

To further explore the prevalence of antagonism for specific ORs expressed in these characterized OSNs, we devised a single-cell RNASeq-based OR identification method. Although we did not exhaustively retrieve the ORs of all indole-responsive OSN groups, we identified an indole-sensitive OR gene family that was amenable to *in vitro* characterization. Most ORs in the family were functionally expressed and responded to most of the indole analogues used for OSN characterization, albeit at various ranges of potencies and efficacies. Olfr745 was the only exception, as it showed only very weak responses to indole, though 2-methylindole did reach saturation.

The high-throughput screening of ∼800 volatile compounds against the Olfr743 family allowed us to probe the prevalence of both agonism and antagonism for a much larger chemical space than the original diagnostic ligands. While this library is small compared to those typically found in drug screens (in which compounds can number in the millions), it is large compared to what has previously been reported in the olfactory literature and is approximately 25% as large as all the odorants reportedly described in public databases (∼3100) [8]. We identified a total of 430 putative antagonists across the receptor family, of which 149 were specific to a single receptor, and the rest shared across multiple members. Clustering the pairwise correlations of the agonist and antagonist single-concentration data across the ∼800 test compounds reflected the phylogenetic clustering of receptors, suggesting a link between sequence diversification and functional diversification.

Examining the top five inhibitors for each receptor in dose-response agonism and antagonism experiments provided more precise and accurate measurements of potency and efficacy across the ten ORs. Both the degree of inhibition and OR-specificity were highly variable, in good agreement with the varied effects observed across groups of functionally segregated *ex vivo* OSNs. Compounds not only elicited distinct ranges of activation or inhibition, but also opposite pharmacological outcome, switching from antagonism to agonism between receptors, effectively leading to lack of functional redundancy. This latter conclusion is in agreement with current views of functional diversification observed for OR paralog activation [51, 52].

To further explore the relationship between phylogeny and function, we identified two sources of diversification in the experimental receptor responses. First, as sequence identity diverged, so did the binding specificity. Even phylogenetically closely related ORs, for which the sets of binders were most similar (Figure S8D), exhibited variable specificity, resulting in distinct molecular receptive ranges. Second, we observed an additional source of variability due to the pharmacological outcomes among paralogous ORs that recognized the same ligands but exhibited opposite responses (i.e. agonism vs. antagonism – Figure S8D). Such distinction upon binding can arise through conformational changes that alter G protein affinity and selectivity, or transduction cascade efficiency [53-55]. This is also observed in OR allelic variability where minimal sequence alterations, often a single SNP, lay outside of presumptive OR binding pockets [56], yet were responsible for dramatic shifts in pharmacological responses [21-23, 27, 28, 57]. Because the OR repertoire evolved along with speciation [58], this functional diversification, and the extra degree of freedom contributed by antagonism, likely contributes to the OR neofunctionalization required for functional gene retention following OR gene duplication.

Our results suggest that antagonism is a prevalent feature of the peripheral olfactory system, and that its combinatorial nature mirrors that of agonism. We demonstrate extensive non-linear OR-mediated computation of mixture information prior to transmission of signal to the olfactory bulb. This was observed directly in calcium imaging experiments for two-component mixtures. The *in vitro* dose-response data across all tested ligands and ORs was shown to be compatible with published models of competitive binding [29, 48-50] and consistent with the theoretically proposed case in which binding affinity was only weakly correlated with activation efficacy. The consequence is to increase the encoding capacity of the system by resisting OR ensemble saturation in the face of complex mixtures of odorants. This and the particular correlation structure of agonistic and antagonistic receptive ranges leads to a far richer mixture encoding logic for the system than one where antagonism is rare and responses are largely additive.

## ACKNOWLEDGEMENTS

We would like to thank the members of Firmenich R&D for their thoughtful comments and advice. In particular, we would like to thank M. Reiter and J. Coulomb for chemical synthesis, J. Pika, M. Emberger and D. Hossain for compound analysis and management.

## AUTHOR CONTRIBUTIONS

Conceptualization, P.P., B.C.S., J.H.B. and M.E.R.; Methodology, P.P., B.C.S., B.J.E., J.H.B., C.T., M.S., R.H.A., G.R., D.A.R. and Z.P.; Formal Analysis, B.C.S., C.T. and G.R.; Investigation, B.J.E., J.H.B., C.T., M.S., R.H.A., G.R., D.A.R., and Z.P.; Resources H.Y.J.; Writing – Original Draft, P.P., B.C.S., B.J.E., J.H.B. and G.R.; Writing – Review & Editing, P.P., B.C.S., B.J.E., J.H.B., C.T., G.R., M.V., and M.E.R.; Visualization B.C.S., J.H.B., B.J.E., C.T., G.R. and D.A.R.; Supervision, M.E.R and M.V.; Project Administration P.P., B.C.S, J.H.B., C.T., M.V.; Funding Acquisition, M.E.R., M.V.

## METHOD DETAILS

### Animal Care & Sources

C57BL/6J male mice, aged 8 to 12 weeks, were obtained from the Jackson Laboratory. All experimental procedures were in compliance with NIH guidelines and were approved by the Mispro Biotech Services Institutional Animal Care and Use Committee.

### Calcium Imaging

Tissue was prepared as described in Poivet et al., 2018 [59]. In brief, olfactory epithelia were placed into 5 mL L15 medium supplemented with 10% fetal bovine serum, 100 U/ml penicillin and 100 μg/ml streptomycin (Gibco BRL, Grand Island, NY, USA), 0.5 U ml−1 collagenase, 1 U ml−1 dispase (Worthington Biochem, Lakewood, NJ, USA), 3.75 mM CaCl2 (Sigma-Aldrich, St-Louis, MO, USA), and 50 μg ml−1 deoxyribonuclease II (Worthington Biochem).

The tissue was incubated at 37°C for 75 min on a rocker, subsequently dissociated by trituration with a siliconized pipette, and plated onto concanavalin-coated glass coverslips (Sigma-Aldrich, 10 mg/mL) placed in 35 mm Petri dishes. Following plating for 30 min to permit cell adhesion, 2 mL of culture medium was added to each dish and the dishes were held at 37°C for at least 1 h. Culture medium consisted of DMEM/F12 (Gibco BRL) supplemented with 10% fetal bovine serum, 1x insulin-transferrin-selenium (Gibco BRL), 100 U/ml penicillin and 100 μg/ml streptomycin (Gibco BRL), and 100 μM ascorbic acid (Sigma-Aldrich).

Cells were loaded with Fura-2 AM (5 μM; ThermoFisher, Waltham, MA, USA; 31°C for 45 min). After being washed with Hank’s Balanced Salt Solution, the coverslips were mounted into a recording chamber. Imaging was carried out at room temperature on an IX83 inverted fluorescence microscope (Olympus, Center Valley, PA, USA) equipped with an Orca-R2 camera (C10600, Hamamatsu Photonics, Hamamatsu, Japan), Proscan III motorized stage (Prior Scientific, Rockland, MA, USA) a Lambda XL light source (Sutter Instrument, Novato, CA, USA), and Lamba-10B optical filter changer (Sutter Instrument). Odorants (Firmenich) were prepared as 1 mM stocks and diluted to a final working concentration in HBSS (recipe), and bath applied using an Agilent 1100 series HPLC system (Agilent Technologies, Santa Clara, CA, USA). A final stimulation with 40 μM Forskolin (Sigma-Aldrich; prepared as 40 mM in DMSO and diluted in HBSS) was made to assess the viability of dissociated OSNs. Images were taken with Metamorph Premier software (Molecular Devices LLC, San Jose, CA, USA) at 340 and 380 nm excitation and 510 nm emission, approximately every 6 s, and there was a 3 min delay between odor stimulation.

### Single Cell RNA-Seq

Single cell picking was done with borosilicate pipettes pulled to a tip opening diameter of 25 ± 5 μm (Cat # B150-86-10, Sutter Instrument) attached to a micromanipulator on microscope stage. Individual cells were expelled into an RNAse-free 200 μl tube and cDNA was generated according to the SMART-Seq v3 Ultra Low Input RNA Kit for Sequencing (Takara, Cat. #634851) for RNA extraction and amplification by 18 PCR cycles. Amplified cDNAs were purified using magnetic bead separation agent Agencourt AMPure XP (Beckman, Cat # A63880) for HiSeq compatible cDNA library generation according to TruSeq RNA Sample Preparation v2 (Illumina, Cat # 15026495). Concentration and integrity of both the cDNA and the libraries were monitored by PicoGreen analyses (BioAnalyzer 2100, Agilent). Illumina HiSeq 100bp single end reads was performed and the resulting reads were first assembled and mapped onto the mouse genome (version mm10, UCSC) and compared to the *ab initio* transcripts (obtained from GENEBUILD) to identify expressed genes. The raw counts were normalized and presented as TPMs (Transcript-Per-Millions).

### Cell line generation

A modified HEK293T cell line expressing the endogenous Rtp1 gene was generated for functional OR expression. A targeted insertion of the constitutively active CMV promoter (P_CMV_) was performed using CRISPR/Cas9 technologies followed by homologous directed DNA repair (HDR). A guide RNA (gRNA) was designed to target positions 148 bp and 147 bp upstream of the Rtp1 gene translation start site and induce a double-stranded DNA break, when combined with Cas9. Two oligo nucleotides (top and bottom target sequence strand) with 3’ overhangs were annealed and cloned into the vector the GeneArt® CRISPR Nuclease Vector Kit to generate the gRNA/Cas9 nuclease plasmid. 5’ and 3’ homology arms (amplified directly from HEK293T genomic DNA, see reagents for primer sequences) flanking the P_CMV_ sequence attached to a puromycin resistance gene (from pPUR vector) were cloned into pGEM®-T Easy Vector to generate the HDR plasmid. HEK293T cells were transfected with 1:1 mixture of gRNA/Cas9 and HDR plasmids using Lipofectamine 2000. After selection in puromycin containing media, single colonies were isolated and tested for functional OR expression. A final clonal cell line was selected for use in this study.

### cAMP Functional Cell-Based Assay

All OR genes were synthesized and cloned into the modified expression vector pME18S-FL containing an SV40 promoter followed by a Flag-Rho tag at the N terminal end of the OR. Expression vectors were transfected at 5μg of DNA and 10μl Lipofectamine 2000 (Life Technologies, Cat. #11668) in 6 ml DMEM per 120 wells into the modified HEK293T cell line, expressing the endogenous Rtp1 gene. Cells were seeded at a density of 7500 cells/well in a volume of 50 µl in 96-well white, opaque bottom plates (Corning, Cat # 3688) in DMEM supplemented with 10% FBS with no antibiotic. Plates were incubated at 37°C in 5% CO2 overnight. Prior to the assay, cells were washed with 100 μl PBS, and incubated with 40 μl of compound solubilized to the correct concentration in DMSO was diluted in each well to a final concentration of 0.3% DMSO in assay buffer (HBSS, 10 mM MgCl2, 20 mM HEPES, 2 mM CaCl2, 0.5 M IBMX). For dose-response agonist experiments, compounds were diluted to 12 different concentrations between 10^-8^ and 10^-3^ M at approximately half-log intervals as above. For antagonist experiments, the EC_80_ of indole was first determined in a duplicate dose-response agonist experiment and then test compounds were diluted as before and mixed with a dilution of indole to achieve the EC_80_. In dose-response experiments in which both agonist and antagonist were varied, compounds were prepared as for the antagonist dose-response experiment but with indole diluted to range between 10^-8^ and 10^-3^ M at half-log intervals as well. Cells were incubated with compound at 37°C in 5% CO2 for 30 minutes In the context of Golf, OR activity was monitored using the HTRF cAMP dynamic 2 kit (Cisbio, Cat # 62AM4PEB), a competitive immunoassay between native cAMP produced by cells and the cAMP tracer molecule, labeled with proprietary CisBio fluorophore D2. cAMP-D2 bound to the anti-cAMP mouse antibody labeled with cryptate generates fluorescence. The signal is expressed as fluorescence ratio of the emission wavelength at 665 nm to 620 nm. The HTRF assay plates were read on Pherastar (BMG labtech). Dose-response data were graphed in GraphPad Prism 7.04 to calculate EC_50_, relative IC_50_, Hill slopes, maxima and minima of the response signals, and to graph dose-response curves. Agonist dose-response experiments were carried out in duplicates. The standard errors of the mean were calculated and the data points were fitted to a four parameter non-linear regression following the Hill equation (unconstrained Hill slope).

### High-Throughput Screening

The same transfection and analyses conditions were applied to the library screening as for the OR full dose-response curve described above. A library of 800 odorants sourced internally was assembled in 100% solvent (DMSO) at a stock concentration of 300 mM and stored at −20° C. For the activation assay, compounds were diluted to a final concentration of 300 μM (DMSO concentration 0.1% final) in a 96 well plate format and delivered to the transfected cells. For the inhibition assay, an initial dose response to indole was performed in triplicate for each receptor to determine the EC_80_ used in each subsequent single point modulation screen. Compounds were blended in a 96 well plate format to a final concentration of 300 μM with indole EC_80_ at 0.2% DMSO final (80 compounds per plate). Single point activity measures were performed for 800 non-redundant binary mixtures and compared to the activity elicited by indole EC_80_ alone. Assay variability and reliability were evaluated by calculating the Z’, a measure of the assay window and the standard deviations of minimum and maximum cAMP production per plate. The average Z’ value obtained across all experiments was 0.86 (SD=0.03) and 0.70 (SD=0.09) for agonism and antagonism respectively, and surpassed minimal requirement above the standard quality limit of Z’ > 0.5 [60]. 

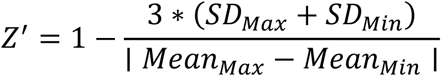

Receiver Operator Characteristic curves were further used to assess the accuracy of the single point screen to predict modulation at the dose-response level. All data points were considered for the Pearson correlation matrix analyses.

#### Competitive binding model with spontaneous receptor activity

We propose a model of competitive binding that includes spontaneous activation of the receptor as shown in Figure 7A. The on and off rates of the binding of ligand X_i_ are denoted by κ^+^_1,i_ and κ^-^_1,i_ respectively. Once bound, there is a rate of activation of the receptor, which depends on the bound ligand. If X_i_ is the bound ligand, we denote its rate of activating the receptor as κ_2,i_. The activated receptor reverts back to its native, unbound state with a rate k^-^_s,_ which can in turn spontaneously activate with a rate k^+^_s_. The output of the experiment is assumed to be proportional to the total number of activated receptors. When the set of ligands X_1_, X_2,_…, X_K_ at concentrations C_1_, C_2,_…, C_K_ respectively are delivered, by writing down the set of coupled rate equations, we can calculate the total number of activated receptors of a particular type at steady state as 

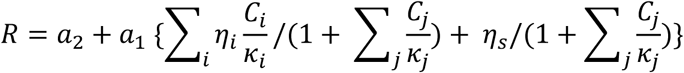

where the sums are from 1 to K, κ^-1^_i_ and η _i_ are the binding affinity and activation efficacy respectively of ligand X_i,_ η_s_ is the constitutive activity of the receptor and *a*_*1*_ and *a*_*2*_ are constants. In terms of the rate constants introduced previously, we have 

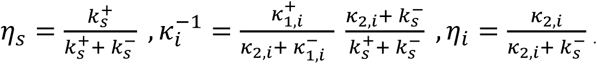

Note that the definitions of η_i_ and η_s_ restrict their range between 0 and 1. If there were no ligands present, all C_i_’s would be zero and the activity is then proportional to η_s_. At large concentrations, all receptors are bound, and all activity is due to ligand-induced activation rather than spontaneous activity. The constants *a*_*1*_ and *a*_*2*_ are independent of receptor-ligand interactions, and depend on cellular processes downstream of receptor activation, number of receptors, reporter properties and other factors.

### Fitting activation and inhibition profiles to the competitive binding model

To obtain the plots in Figure 7B, we first normalized the data between zero and one, where zero corresponds to the HTRF ratio of the lowest concentration of the activator (0.1μM) and no inhibitor and one corresponds to the saturation level of the activator dose-response profile with no inhibitor. Performing this normalization is equivalent to subtracting *a*_*2*_ *+ a*_*1*_η_s_ and then dividing by *a*_*1*_(η_act_ - η_s_) from the equation for *R* from the previous Methods section. Here η_act_ is the unknown activation efficacy of the activator. If η_inh_ is the activation efficacy of the inhibitor, we can write the normalized response in terms of the concentrations of the activator and inhibitor and their binding affinities as 

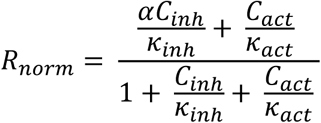

where α is the ratio (η_inh_ - η_s_)/(η_act_ - η_s_). Note there are three free parameters: κ_inh,_ κ_act_ and α. We obtain the best fit curves in Figure 7B by minimizing the RMSE between the CB model and the data from the competitive binding assay.

To obtain the values of the binding affinities in Figure 7C and D, we fit the CB model to the activation and inhibition profiles of 39 unique compounds delivered to the ten receptors from the Olfr743 family. For the activation assay, we first subtracted the HTRF ratio at the lowest concentration of the activator, which is equivalent to subtracting *a*_*2*_ *+ a*_*1*_η_s_ from the equation for *R*. We fit the dose-response curves to the Hill function with Hill coefficient 1 (as predicted by the CB model). We collected the best-fit binding affinities for the receptor-ligand pair which show a significant positive activation of the receptor (std. dev of the HTRF ratio across the ten concentrations > 80 and positive best-fit saturation level) while excluding those which have a best-fit ln κ^-1^ > 8.5 ∼ 200μM (since the tested concentrations do not exceed much beyond this value). The screening above yielded the binding affinities for 50 activators from a total of ∼400 activation profiles. For the inhibition assay, the activator was indole delivered at EC_80_, calibrated from an activation assay performed on the same set of cells. We normalize the data similar to the analysis from the competitive binding assay. The normalized inhibition profiles are fit to the equation for *R*_*norm*_ given above, where κ_act_ is obtained from the activation assay for the activator and *C*_*act*_ is the EC_80_. The best-fit binding affinities which show significant inhibition (std. dev of the HTRF ratio across the ten concentrations > 80, best-fit α value < 0.5, and best-fit ln κ^-1^_inh_ > 8.5) were collected, yielding the binding affinities for 149 inhibitors.

Note that the best-fit curves from the CB model yield the activation efficacy η in a continuum between 0 and 1. In practice, however, most profiles do not saturate and the best-fit η value is imprecise. In our dataset, we observed few partial agonists and thus we simplified our analysis by effectively projecting the activation efficacies into two broad categories: activators (η ≈ 1) and inhibitors (η ≈ 0).

### Correlation between binding affinity and functional outcome

We calculated the Pearson correlation coefficient between the logarithm of the binding affinity and the binary variable activator/inhibitor. Using the expression for the correlation coefficient between a continuous and binary variable, we have 

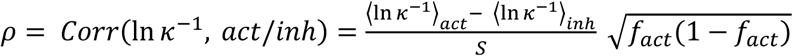

where *S* is the empirical standard deviation of ln κ^-1^ (= 1.37), and *f*_*act*_ is the probability that a compound that binds to a receptor is an activator. Using the data from the high-throughput screen of 800 odorants we estimated *f*_*act ∼*_ 0.68 (444 total activators out of 653 total binders), which gives ρ = 0.18. To calculate the standard deviation of ρ, we repeat the entire analysis by first adding 6% noise to the dataset, where 6% is the average noise to signal ratio estimated from measurements of the HTRF ratio at the lowest concentration of each compound. The standard deviation of ρ is then computed to be the standard deviation of the re-calculated ρ values over 100 repetitions. Note that the correlation coefficient calculated here corresponds to the correlation between log-binding affinity and activation efficacy when the odorant and receptor are both independently drawn in each sample. One could instead calculate the correlation coefficient for each receptor, where the receptor is fixed and the correlation is calculated for sampled odorants. The latter would indeed contain more information about each receptor, but requires enough data to compute ρ for each receptor, which is not available with our current dataset.

To generate the plots in Figure 7E, we calculate the probability that a receptor is activated when an odorant mixture consisting of a particular number of ligands was delivered. We assume that the receptor-ligand interactions are independent across ligands, that if a ligand binds, it is either a perfect agonist (η = 1) or a perfect inhibitor (η = 0) and that each ligand is equiproportionate and delivered at saturating concentrations (the normalization effect is even stronger at weaker, unequal concentrations, see ref. 29). First, we assume the fraction of ligands that bind to a receptor (i.e., they either activate or inhibit the receptor) to be ∼30%, as estimated from the data. Second, if the ligand binds, it is an activator with probability *f*_*act*_ = 0.68. The logarithm of the binding affinity of an activator (inhibitor) is drawn from a normal distribution with mean *M*_*act*_ (*M*_*inh*_) and standard deviation 1.37. *M*_*act*_ = 11.2 and *M*_*inh*_ = 10.62 correspond to the experimentally obtained values, giving ρ = 0.18. From the CB model, the activity of the receptor in the presence of a saturating, equiproportionate mixture is 

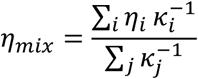

where the sum is over the ligands in the mixture. If η_mix_ > *f*_*act*_, the receptor is considered active. The sparsity for a fixed mixture complexity K is defined as the probability that η_mix_ > *f*_*act*_ over many samples of mixtures with K ligands. To obtain the sparsity vs K for other values of ρ as shown in Figure 7E, we tune the value of *M*_*act*_ accordingly.

## QUANTIFICATION AND STATISTICAL ANALYSIS

### Calcium Imaging Analysis

Images collected during the experimental phase were analyzed by a custom protocol built in Pipeline Pilot (BioVia Dassault Systèmes, San Diego, CA). Each image was tagged with a timepoint during acquisition, which could then be matched to the time of odorant administration. The frameset corresponding to the positive control (Forskolin) injection was isolated for image segmentation. A ratio was created of the 340 nm/380 nm image pair for each timepoint, followed by the creation of a 3D stack of all the resulting images. This 3D image stack was projected into a 2-dimensional image where each pixel is equal to the 85^th^ percentile order statistic of that pixel location in the original 3D stack. Cell regions were then determined using adaptive thresholding of 15% above the mean for a window size of 16×16 pixels. The center of intensity peak for each region fed into a watershed segmentation to split clusters into individual cells. These individual cells were then filtered on contour eccentricity, creating a final segmentation image.

The ratios of the 340 nm/380 nm images for the full experiment were then calculated, and the mean region intensities at every timepoint for each cell defined by the segmentation image was measured and scaled by a factor of 100. The background (defined by the average intensity of the 5 frame prior to the subsequent injection) was then subtracted for each injection frame to give the delta values. Peak characteristics (slopes of peak and post-peak, fluctuation, post-peak upward deflection, area under the curve, and signal-to-noise ratio) of each cell for each injection window were fed into a trained Random Forests model to predict a PEAKTAG (1, 0, −1) relating to good, flat, or bad peaks, respectively. All injection frames for a given cell were then merged to give each cell a ‘good’ or ‘bad’ call based on PEAKTAGs, a delta value of at least 20 for the Positive control frame, and a PEAKTAG of 0 during the Negative control frame. A figure of merit (FOM) was calculated by the accumulation of bad peaks over the total number of injections per cell. All cells with a FOM greater than 0.0 were validated by hand to ensure data fidelity.

To estimate the number of unique odor responsive groups, the odorant responses by each cell were arranged into a binary array such that 0 indicated no response, while 1 represented a validated response to that odorant. Any cells that responded inconsistently to the same odorant were rejected from further analysis. The pattern of activation for each cell allowed for classification into distinct groups. All such cell activation patterns were fed into a classifier in a randomized order. The emergence of new groups and the growth of previously seen groups were tracked and fed through a rarefaction analysis protocol (iNEXT R package,[61]). This allowed for the estimation of the expected total amount of groups, based on the abundance and growth rates of the already identified activation patterns.

Cells that have been validated as “good cells” were fed into a Modulation Analysis tool, custom built in Pipeline Pilot. Any cells that responded to stimuli inconsistently (i.e. cells with spurious peaks) were rejected. Each cell that passed had a linear regression calculated to compensate for the natural attrition seen over the course of an experiment. A best fit line was drawn for the peaks corresponding to the agonist of interest. Cells with a slope that would cause an interception with the baseline before the Forskolin injection were rejected. The modulation value was calculated as the percent difference from the best fit line at that position. A negative modulation value indicates a reduction in the response when compared to the agonist of interest, and a positive modulation value shows and increased response. Where Δf is the maximum peak height of the agonist and potential antagonist mixture, m is the slope of the best fit line, InjFr is the time point of the mixture injection, and b is the intercept as determined by the best fit line, the modulation value can be determined as: 

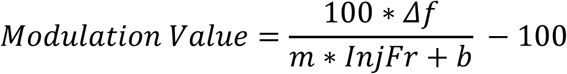

### Statistical Analyses

Figures were assembled with GraphPad Prism, Inkscape and RStudio software. Phylogenetic analyses were performed in BioEdit with the Protdist v3.5c application. Violin plots were made with Vioplot [62] and Tidyverse [63] R packages. Rarefaction analyses were carried out using iNEXT R package [61].

## DATA AND SOFTWARE AVAILABILITY

Requests for raw data and instrumentation and analysis code should be directed to the Corresponding Author.

---

**Figure S1.**
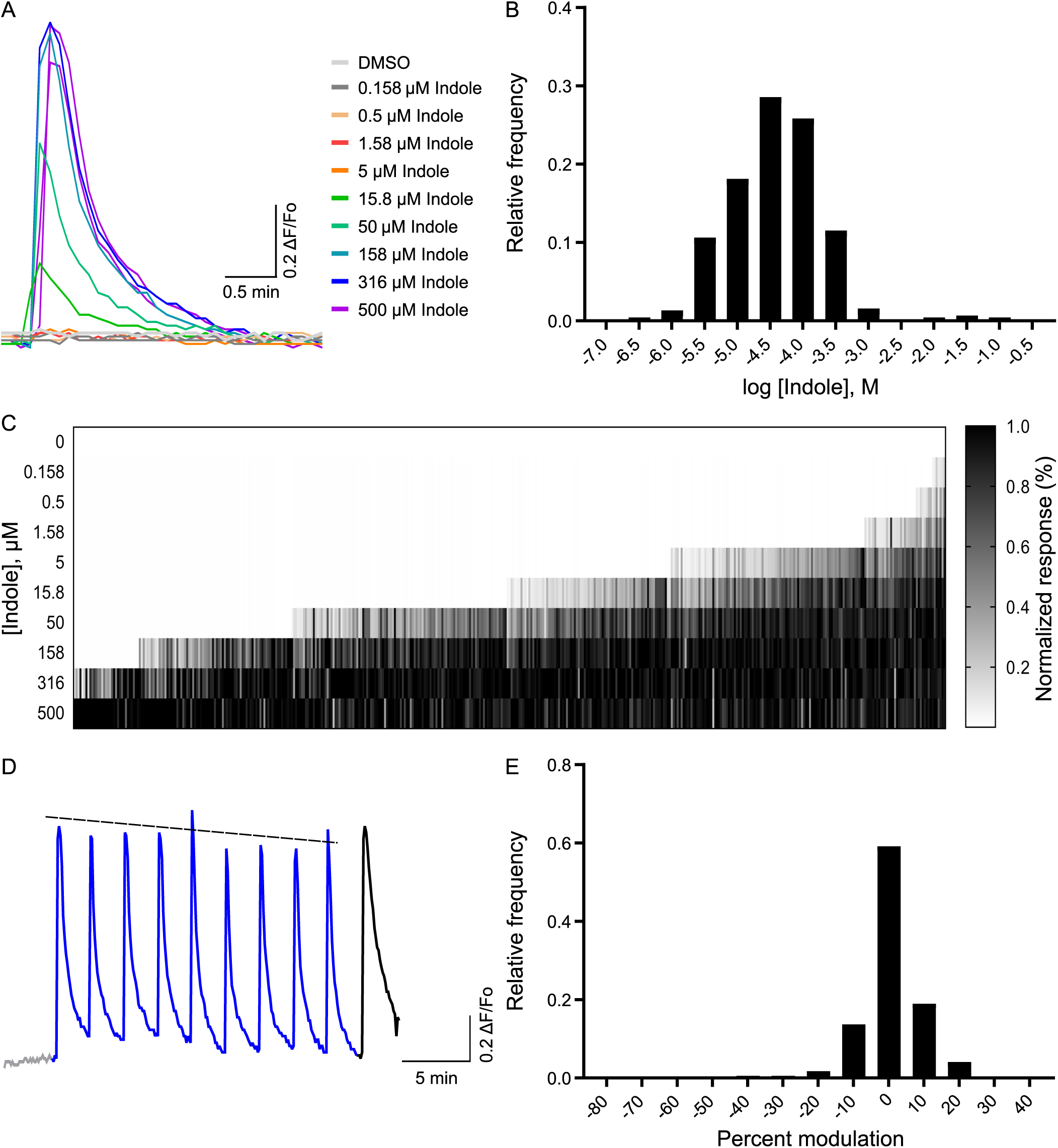
Indole activated dissociated olfactory sensory neurons in a concentration-dependent manner. (A) An example of an OSN calcium-imaging response to increasing concentrations of indole (ranging from 0.158 to 500 µM). (B) Population histogram of the distribution of EC50s for OSNs responding to indole. (C) Population heatmap showing the concentration-dependent recruitment of indole-responsive OSNs. Each column represents the normalized response of a single OSN to increasing concentrations of indole. The lowest and highest concentrations of indole were repeatedly administered to verify responses; cells with spurious responses were discarded. Response strength was normalized to the odor peak response in a given cell. For B and C, 441 OSNs from N = 3 experiments. (D) An example of an OSN response to repeated administration of 25 µM indole (blue), 0.3% DMSO (negative control; gray), and 40 µM Forskolin (positive control; black). (E) Population histogram of the distribution of percent modulation observed between alternating presentations of indole in D.

**Figure S2.**
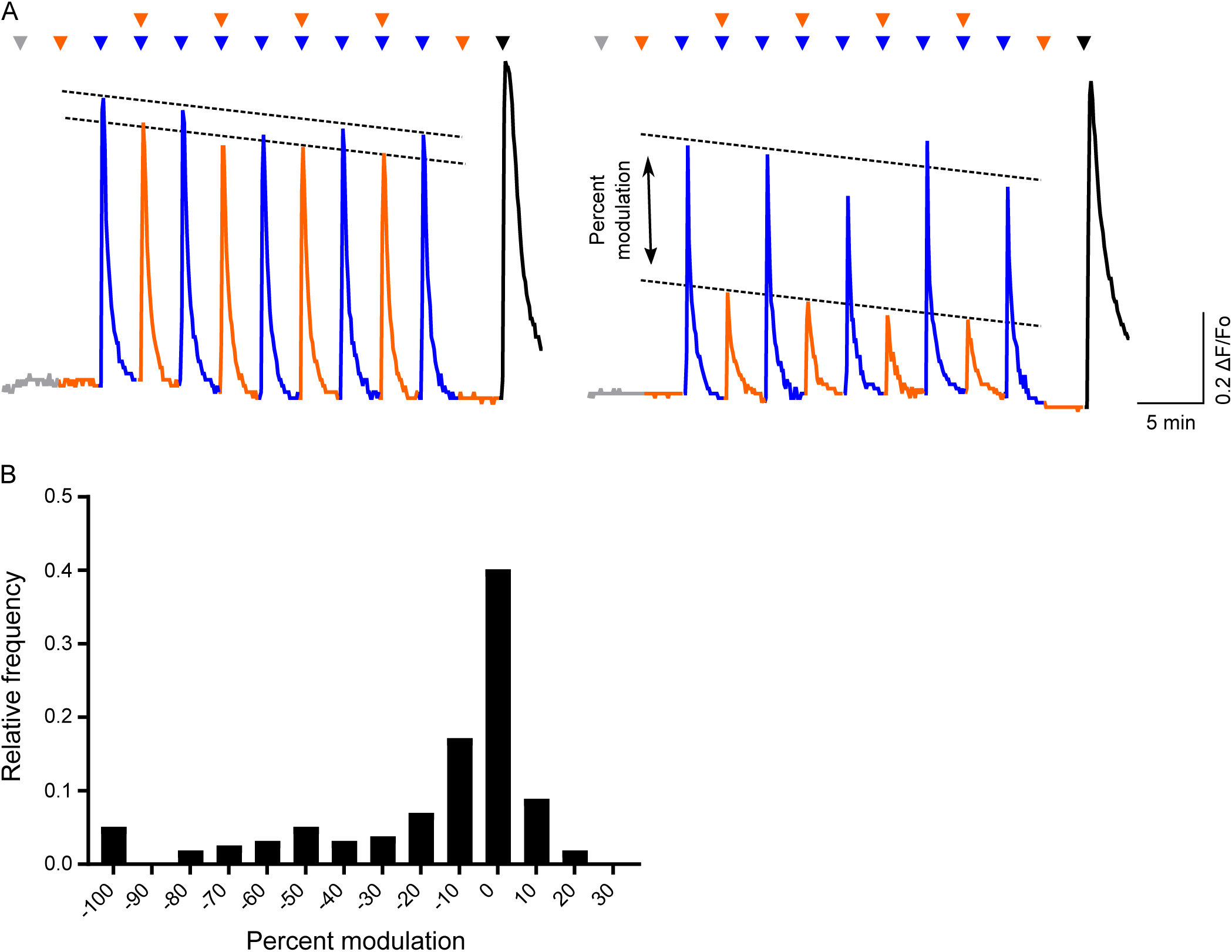
Experimental design for modulation test in dissociated OSNs. (A) Calcium imaging of OSN responses to the repeated administration of 25 µM indole or to a mixture of 25 µM indole and 125 µM Lilyflore^®^, permitting measurement of the percent modulation of the indole response by Lilyflore^®^. Triangles denote the time of administration (gray, 0.3% DMSO, negative control; blue, indole; orange, Lilyflore^®^; black, 40 µM Forskolin, positive control). Forskolin is an activator of adenylyl cyclase [64], and is used as a measure of maximal activation of the signal transduction cascade in OSNs. (B) Population histogram of the Lilyflore^®^ modulation of indole-activated OSNs as described in A (157 OSNs; N = 4).

**Figure S3.**
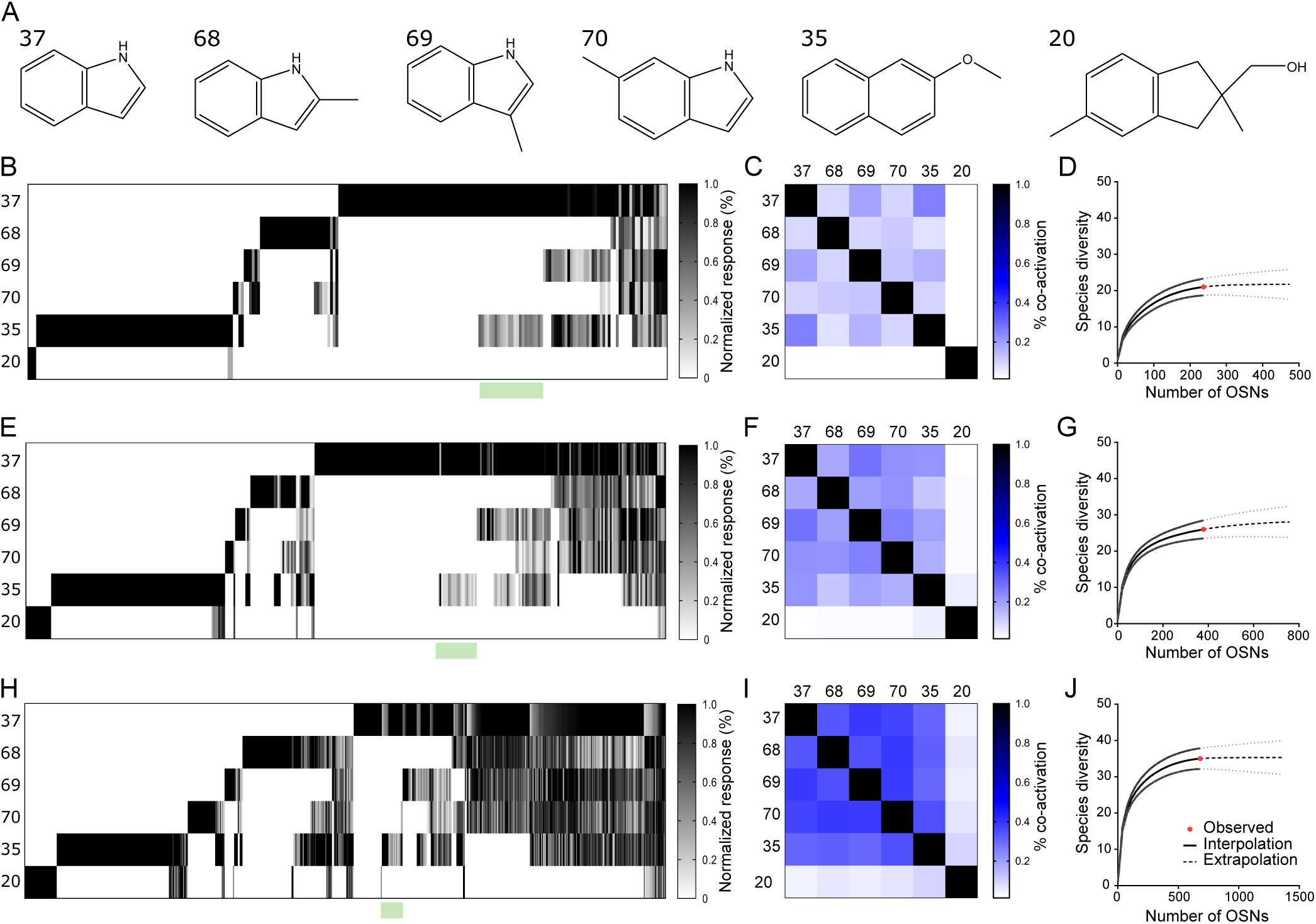
Indole and related compounds activated distinct but partially overlapping populations of OSNs. (A) Chemical structures of indole (37) and five structural analogues including 2-methylindole (68), 3-methylindole (69), 6-methylindole (70), 2-methoxynaphthalene (35), and Lilyflore^®^ (20). (B) Population heatmap of the normalized response of 237 OSNs to the administration of individual odorants in A, where each was presented in duplicate at 5 µM and responses were measured by ratiometric calcium imaging (18,899 Forskolin+ OSNs sampled, N = 3 experiments). Each column represents the average odor-induced activation of a single OSN normalized within cell to the peak odor response. All cells responded to 40 µM Forskolin (data not shown). (C) Response correlation matrix for the six odors in A when presented at 5 µM. (D) Rarefaction analysis was performed to test whether the OSN population size sampled was large enough to likely identify all possible activation profiles at 5 µM. (E-G) Same as in B-D but odorants were delivered at 50 µM (N = 379 OSNs of 19,485 sampled). (H-J) Same as in B-D but odorants were delivered at 250 µM (N = 680 OSNs of 15,013 sampled).

**Figure S4.**
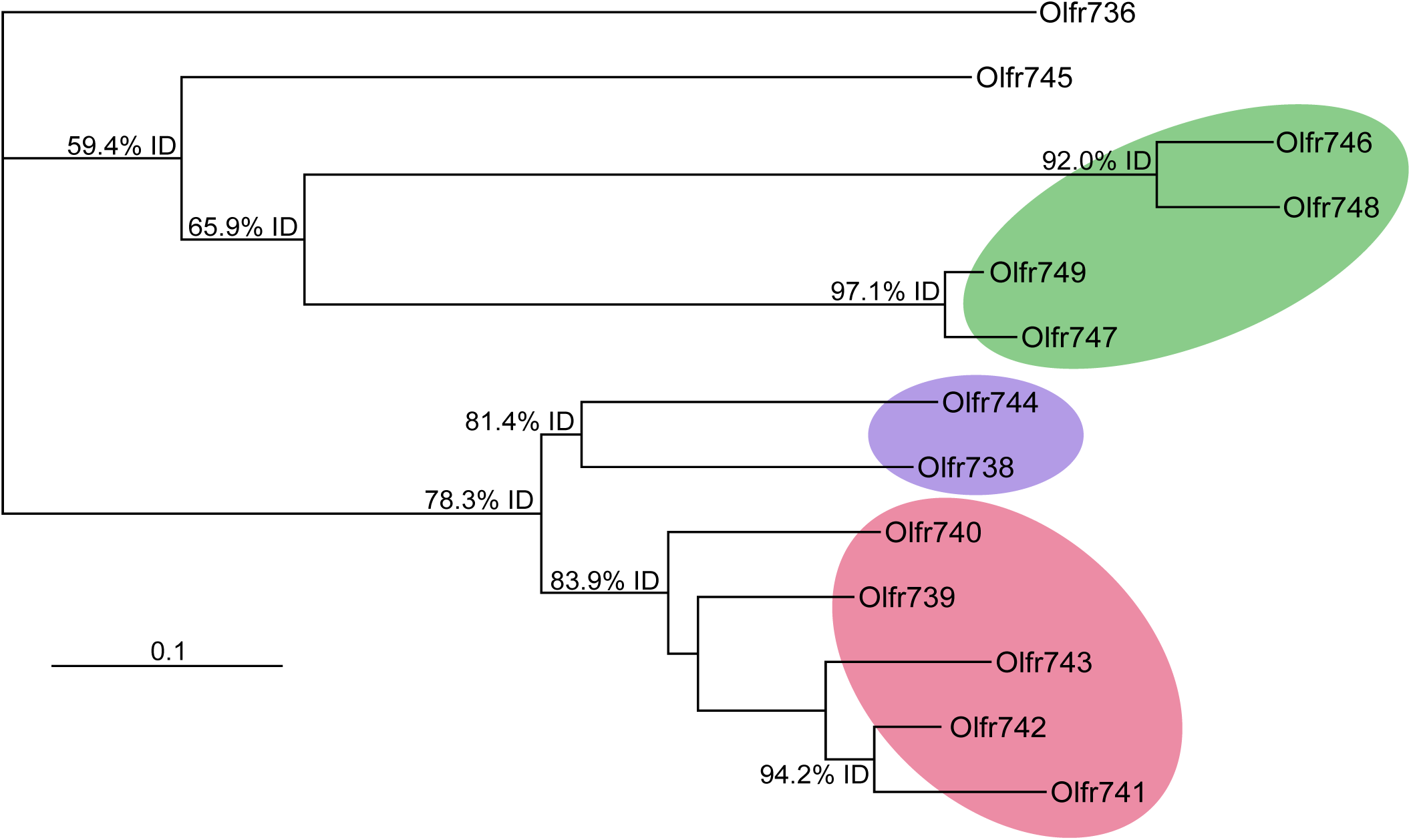
Phylogenetic relationship between 12 paralogs based on an amino acid alignment. Percent identity levels are indicated for select nodes. Olfr736 represents the outgroup and did not elicit a response from any test ligands. Scale bar: average number of substitutions per site

**Figure S5.**
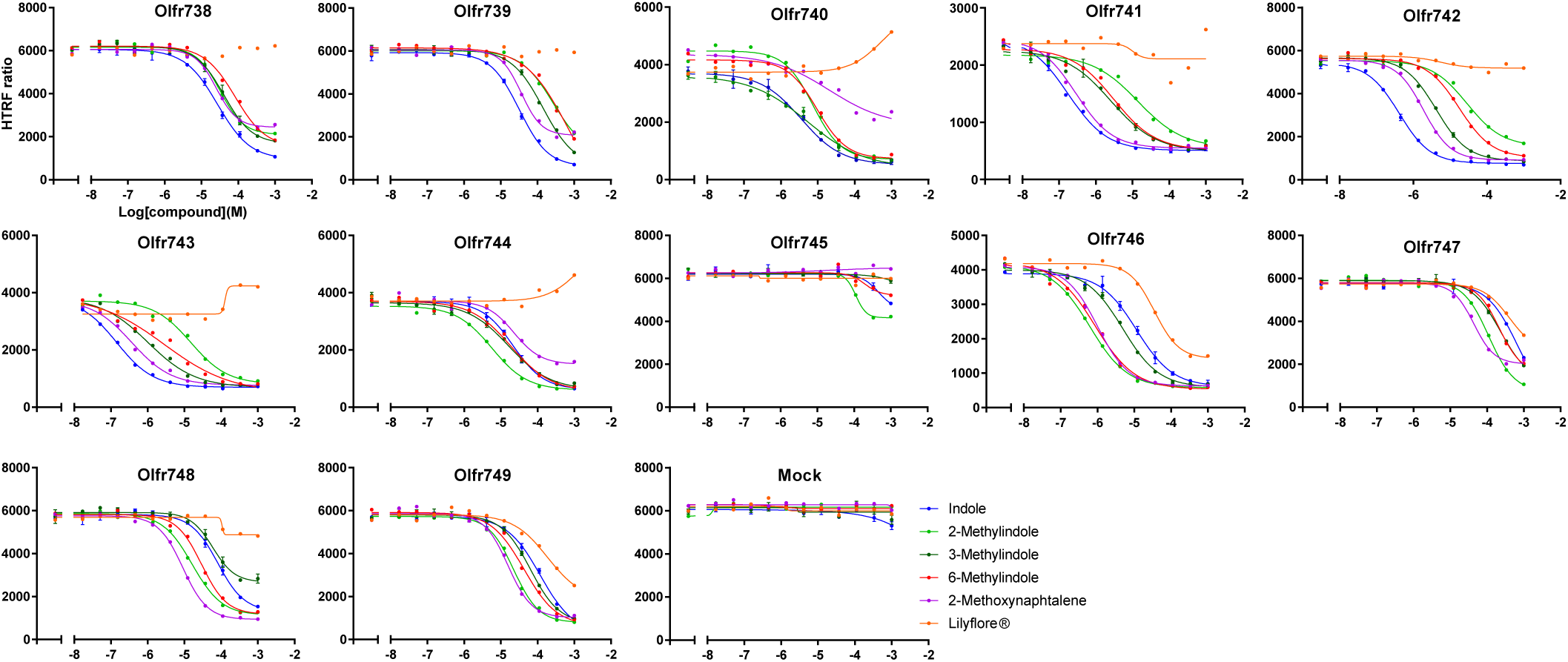
Functional diversity of OR paralogs to indole-derived compounds. Dose-responses of the 12 OR paralogs used to generate the heatmap in Figure 5F. Receptors were probed with indole (dark blue), 2-methylindole (green), 3-methylindole (light blue), 6-methylindole (red), 2-methoxynaphthalene (purple), Lilyflore^®^ (orange). The empty vector control is shown. Both potencies and efficacies varied by receptor and revealed variation in activation levels among phylogenetically related ORs. Lilyflore^®^ activation of indole sensitive ORs only occurred at high concentrations for receptors Olfr746 to Olfr749, consistent with the low OSN response overlap observed in Figure 2 between indole and Lilyflore^®^ at several concentrations

**Figure S6.**
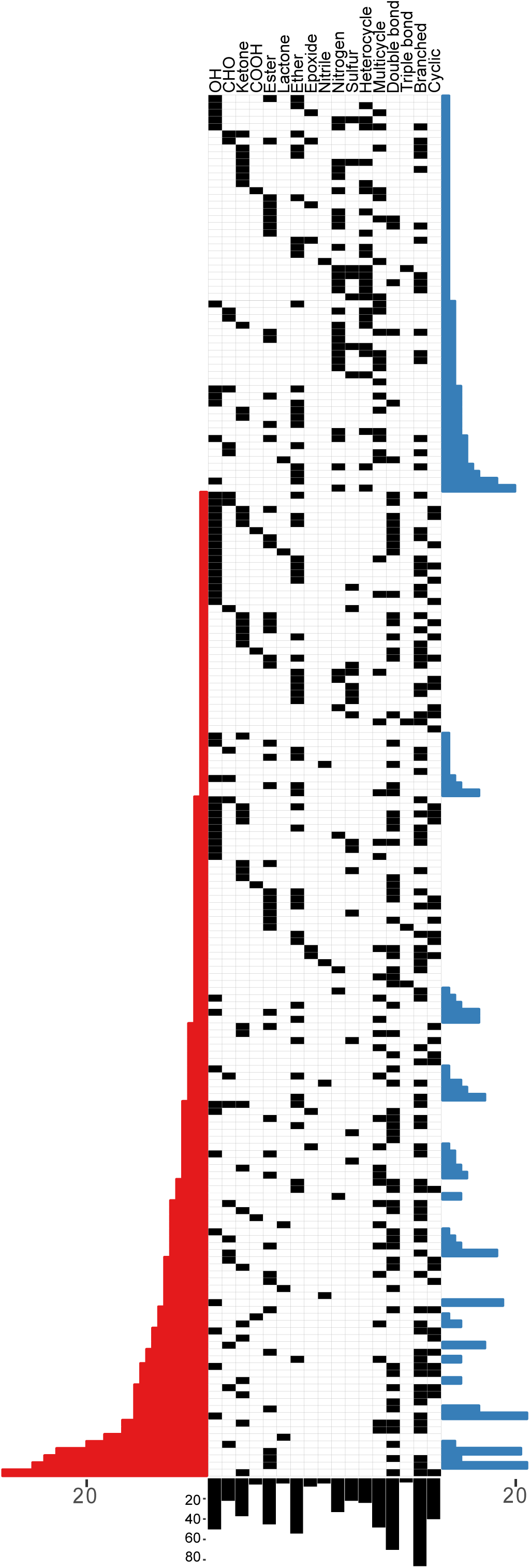
Characterization of screening library in terms of presence of specified chemical features. The chemical diversity of the 806 compounds contained in the high-throughput screening library is represented. Cells in the central matrix indicate the presence (black), or absence (white) of specified chemical features indicated in the column headers. Histograms to left and right show the number of aliphatic and aromatic compounds, respectively. Bars beneath indicate the total number of compounds possessing each named feature.

**Figure S7.**
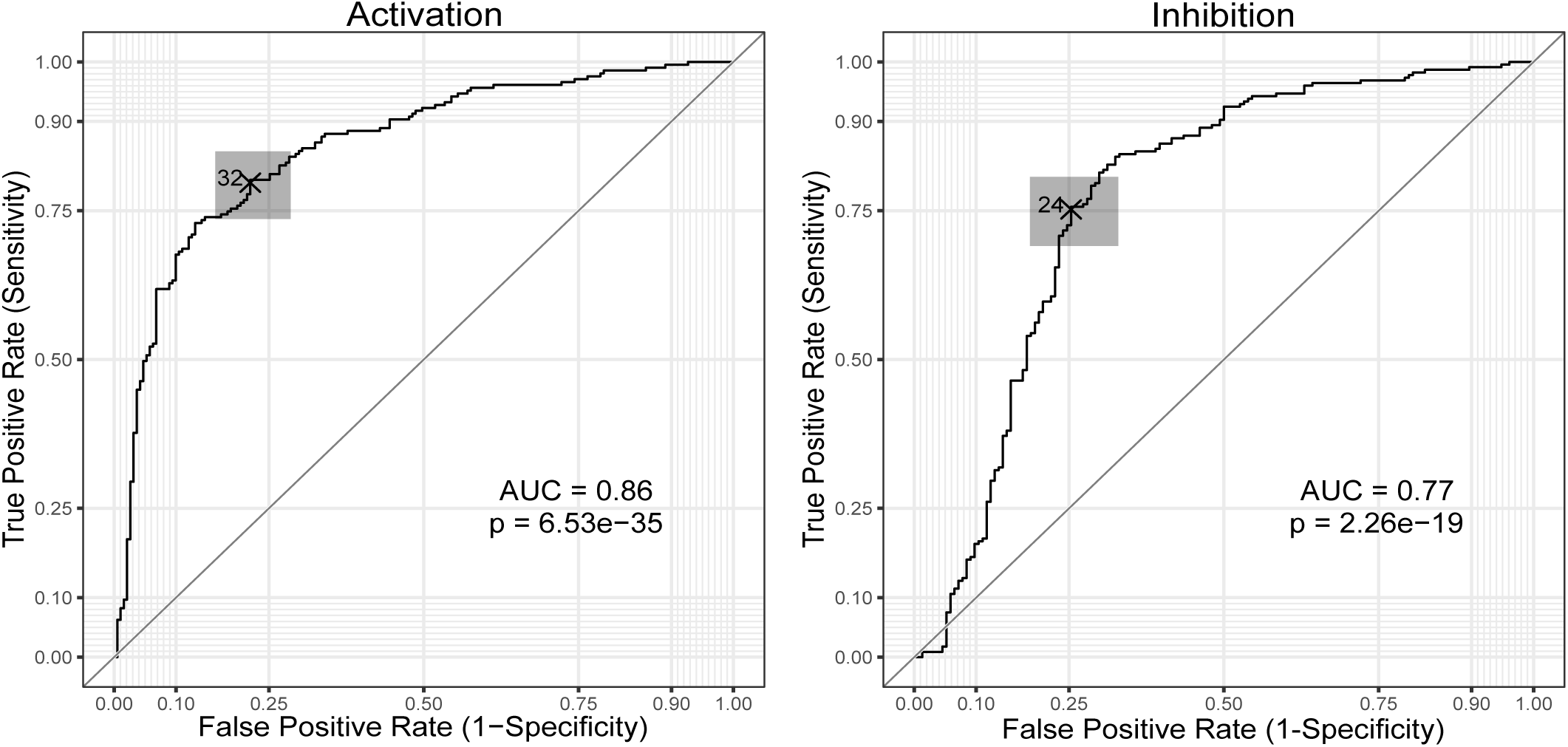
Statistical Receiver Operator Characteristic (ROC) analysis. ROC curves demonstrate that both primary activation and inhibition screens were able to predict whether compounds would pass an activation or inhibition dose response experiment. The optimal threshold, determined as the closest point to the upper left corner, was 32% for activation and 24% for inhibition. Confidence interval for the threshold is shown in gray. AUC = area under the curve.

**Figure S8.**
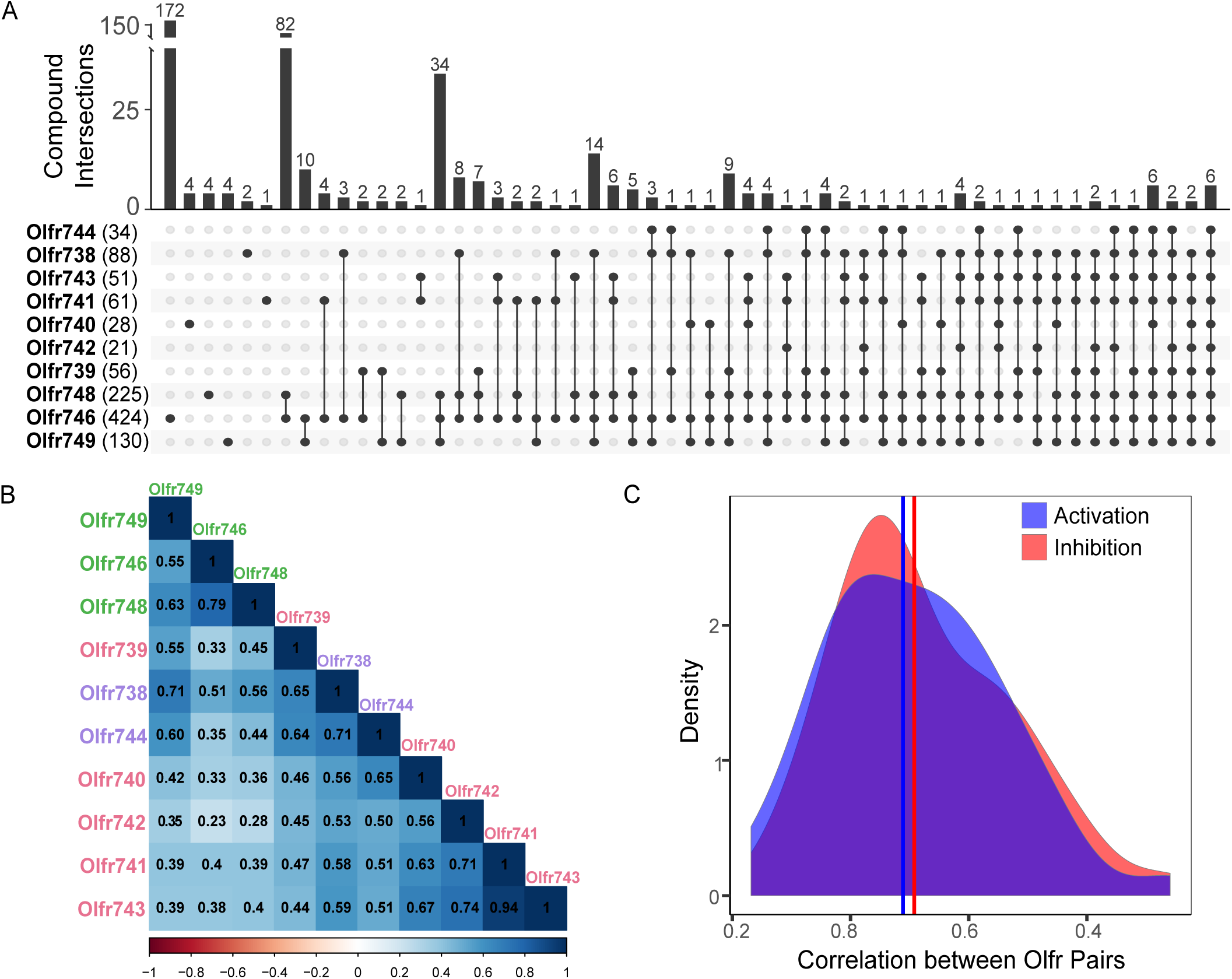
Large inhibition screen reveals agonism diversity among closely related indole-sensitive ORs. (A) Upset plot showing the number (bar height) of activators (percent activation > 32%) which were unique (single dots) or shared (linked dots) among Olfr743 family members. (B) Pearson correlation in activation screening values for each pair of ORs. Label colors indicate phylogenetic clades inferred by amino acid sequence similarity. (C) Distribution of correlation in activation and inhibition screening values for all receptor pairs. Vertical line indicates median correlation for each screen.

**Figure S9.**
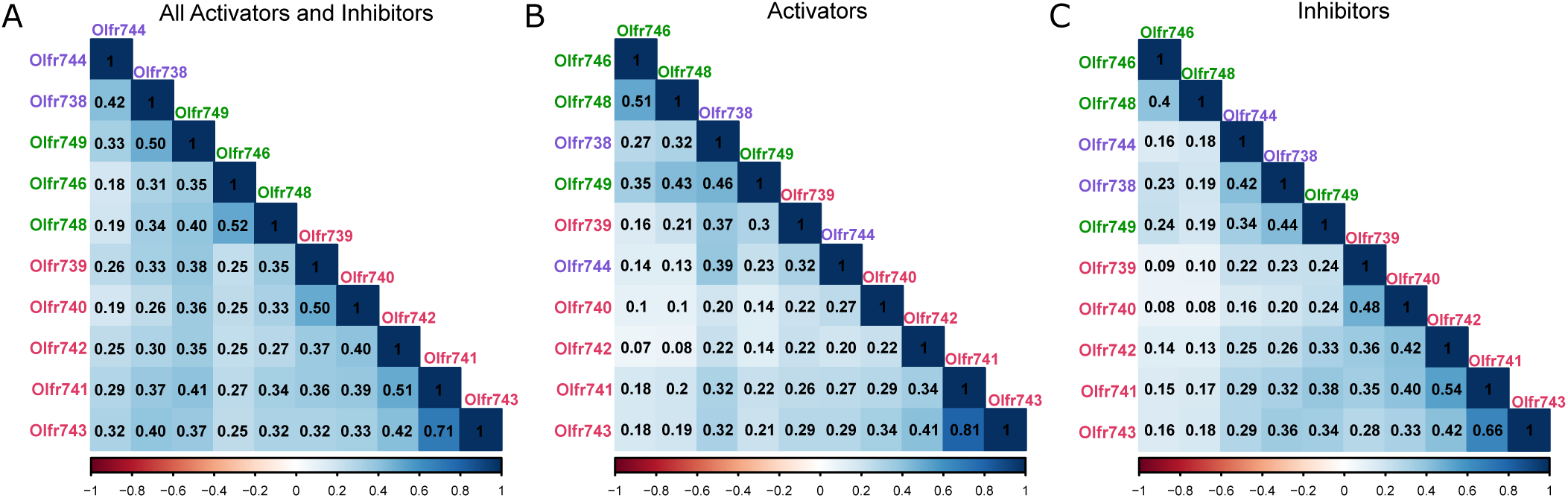
Binding properties conservation between paralog ORs. The Jaccard Indices for (A) activators and inhibitors together, (B) activators only, or (C) inhibitors only, for each pair of ORs is shown. Label colors indicate phylogenetic clades inferred by amino acid sequence similarity. (D) Distribution of Jaccard Indices for all OR pairs for activators and inhibitors together, activators alone, and inhibitors alone. Vertical lines indicate the median Jaccard Index for each condition.

